# Connecting Transcriptomics with Computational Modeling to Reveal Developmental Adaptations in the Human Pediatric Myocardium

**DOI:** 10.1101/2024.04.19.589826

**Authors:** Shatha Salameh, Devon Guerrelli, Jacob A. Miller, Manan Desai, Nicolae Moise, Can Yerebakan, Alisa Bruce, Pranava Sinha, Yves d’Udekem, Seth H. Weinberg, Nikki Gillum Posnack

## Abstract

**Background:** Nearly 1% or 1.3 million babies are born with congenital heart disease (CHD) globally each year – many of whom will require palliative or corrective heart surgery within the first few years of life. A detailed understanding of cardiac maturation can help to expand our knowledge on cardiac diseases that develop during gestation, identify age-appropriate cardiovascular drug therapies, and inform clinical care decisions related to surgical repair, myocardial preservation, or postoperative management. Yet, to date, our knowledge of the temporal changes that cardiomyocytes undergo during postnatal development is largely limited to animal models.

**Methods:** Right atrial tissue samples were collected from n=117 neonatal, infant, and pediatric patients undergoing correct surgery due to (acyanotic) CHD. Patients were stratified into five age groups: neonate (0-30 days), infant (31-364 days), toddler to preschool (1-5 years), school age (6-11 years), and adolescent to young adults (12-32 years). We measured age-dependent adaptations in cardiac gene expression, and used computational modeling to simulate action potential and calcium transients.

**Results:** Enrichment of differentially expressed genes (DEG) was explored, revealing age-dependent changes in several key biological processes (cell cycle, cell division, mitosis), cardiac ion channels, and calcium handling genes. Gene-associated changes in ionic currents exhibited both linear trends and sudden shifts across developmental stages, with changes in calcium handling (*I*_NCX_) and repolarization (*I*_K1_) most strongly associated with an age-dependent decrease in the action potential plateau potential and increase in triangulation, respectively. We also note a shift in repolarization reserve, with lower *I*_Kr_ expression in younger patients, a finding likely tied to the increased amplitude of *I*_Ks_ triggered by elevated sympathetic activation in pediatric patients.

**Conclusion:** This study provides valuable insights into age-dependent changes in human cardiac gene expression and electrophysiology among patients with CHD, shedding light on molecular mechanisms underlying cardiac development and function across different developmental stages.

## Introduction

Nearly 1% or 1.3 million babies are born with congenital heart disease (CHD) globally each year – many of whom will require palliative or corrective heart surgery within the first few years of life^1^. Accordingly, cardiothoracic surgery remains the third-highest volume procedure in the United States among pediatric patients, accounting for over 20% of all surgeries in children under 3 years of age^2^. Younger age at the time of surgery is considered a significant risk factor for adverse postoperative outcomes, including myocardial injury, cardiac arrhythmias, and depressed mechanical function^3–5^. As such, intraoperative injury and/or postoperative recovery can be influenced by the structural, functional, and metabolic immaturity of the pediatric myocardium^6^. A more detailed understanding of cardiac maturation will help to expand our knowledge on cardiac diseases that develop during gestation, identify age-appropriate cardiovascular drug therapies, and inform clinical care decisions related to surgical repair, myocardial preservation, or postoperative management.

During postnatal life, the mammalian heart undergoes a series of developmental changes that facilitate the transition from the intrauterine to extrauterine environment. Throughout this developmental process, cardiomyocytes adjust to hemodynamic demands by increasing their size, enhancing structural organization, and adapting their electrophysiological and contractile properties^7^. Studies suggest that human cardiomyocytes evolve within the first year of life – but, may not reach full maturity until nearly a decade after birth. For example, one hallmark of cardiomyocyte maturation is the transition from hyperplasia (cells proliferate by cell division) to hypertrophic growth (cell size increases without cell division). In humans, the proliferative capacity of atrial cardiomyocytes declines within the first year of life – wherein the percentage of proliferating cardiomyocytes from 3-month-old hearts is 11 times greater than 6-month-old hearts and 27 times greater than 12-month-old hearts^8^. Additional biomarkers of postnatal maturation include sarcomeric protein isoform switching^9–11^, maturation of excitation-contraction coupling^12,13^, age-dependent adaptations in cardiac ion channel expression and function^14–18^, and the assembly of intercalated discs^19^. Regarding the latter, the spatiotemporal organization of key proteins involved in mechanical and electrical intercellular connections appears to be underdeveloped in the human myocardium until ∼7 years of life^19^.

To date, our knowledge of the temporal changes that cardiomyocytes undergo during postnatal development is largely limited to animal models^20,21^ – due in part to the scarcity of human cardiac tissue for use in biomedical research. To overcome this limitation, we collected right atrial tissue samples from patients undergoing open heart surgery. In this study, we first evaluated age-dependent adaptations in atrial gene expression in acyanotic CHD patients (5 days – 32 years old) with normal oxygen saturation levels. We then used an *in silico* approach to translate experimental gene expression measurements to ionic current conductances using an established human atrial specific computational model^22–24^. To the best of our knowledge, this is the first study to couple experimental data with computational modeling to predict action potential and calcium transient morphology throughout pediatric development.

## Methods

Detailed methods are provided in the **Supplemental Material**; data and detailed protocols are available upon reasonable request. Please see **Supplemental Table 1** for patient demographic information, **Supplemental Table 2** for action potential and calcium transient biomarker information, and **Supplemental Table 3** for gene selection based on ionic currents. Human subjects experiments adhered to protocols approved by the Children’s National Hospital Institutional Review Board-approved protocol (IRB#Pro00012146). Transcriptomic data has been deposited to the Gene Expression Omnibus.

## Results

### Differential Gene Expression throughout Postnatal Development

Age-dependent adaptations in cardiac gene expression are documented in animal models^20,21^, but the timing and extent to which these adaptations occur postnatally in humans is unknown. Accordingly, we performed microarray analysis to identify differentiall expressed genes (DEGs) in acyanotic CHD patients and define the biological pathways associated with cardiac development and maturation (**Figure 1**). Using 1.25-fold expression cut-off and a false discovery rate of 0.1, a total of 1321 DEGs were identified in atrial samples collected from neonates versus adolescent/young adults (**Figure 1A,B; Supplemental Table 5**). Of these DEGs, 43% were upregulated and 57% were downregulated in neonates. A comparable trend was observed in infants, whereby 1163 DEGs were identified compared to adolescent/young adults (39% upregulated, 61% downregulated). At older ages, there were a smaller number of DEGs, suggesting that developmental adaptations begin to taper after the first year of life. Of interest, only five DEGs were identified between the two oldest age groups – school aged children versus adolescent/young adults (*VIT, NKAIN2, MBNL3, CPNE8, TEMEM71*). Volcano plots (fold change versus log_10_ p-value) and Venn diagrams were generated to illustrate DEGs that appear to be developmentally regulated (**Figure 1C-E**). As expected, neonates and infants had considerably overlap in the genes that were downregulated (221) and upregulated (110) – compared to the adolescent/young adult group (**Figure 1E**). Collectively, younger patients overexpressed genes related to cell cycle, cytokinesis, and immature sarcomeric protein isoforms (e.g., *MKI67, ANLN, TOP2A, CDK1, TNNI1*) and underexpressed genes involved in contractile function (e.g., *S100A1, NEB, ACTA1, TPM3*).

**Figure 1.**
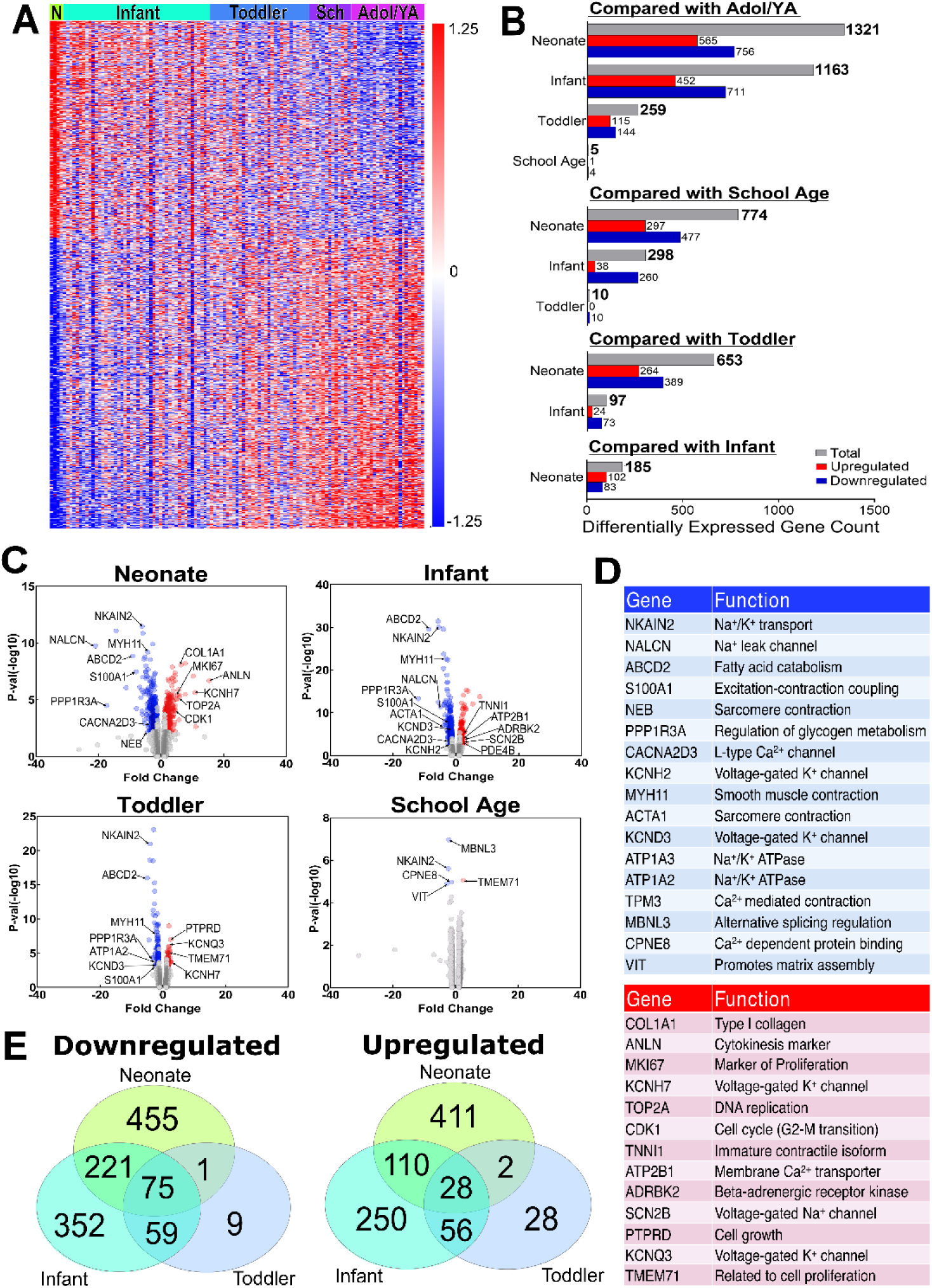
**(A)** Differentially expressed genes were identified for each age group, relative to adolescent/young adult cohort (Adol/YA), using one-way ANOVA with a 1.25-fold change cutoff and a 0.1 false discovery rate. A total of 1321 differentially expressed genes are shown in the heatmap (each row) for each patient (column); on a per gene basis, the signal intensity was log_10_ transformed and Z-score normalized. **(B)** Total number of differentially expressed genes (gray), including those upregulated (red) and downregulated (blue) between each age group. **(C)** Volcano plot illustrates differentially expressed genes that are upregulated (red) or downregulated (blue) for each age group, relative to the adolescent/young adult cohort. Coordinates indicate the fold change relative to the p-value. A subset of genes are labeled, and **(D)** functional description shown. **(E)** Venn diagrams illustrate differentially expressed genes (relative to adolescent/young adults) that are unique or common between the three youngest age groups (neonate, infant, toddler).

### Functional Annotation and Enrichment Analysis

To identify gene ontologies and biological pathways associated with myocardial development and maturation, we used an array of functional annotation and enrichment analysis tools (**Figure 2**). Results of Protein Analysis Through Evolutionary Relationships (PANTHER) classification analysis by biological process are shown in **Figure 2A** (neonate vs adolescent/young adult) and **Figure 2E** (infant vs adolescent/young adult). DEGs were classified similarly in neonates and infants, with the largest proportion of genes connected to cellular process (GO:0009987), biological regulation (GO: 0065007), localization (GO:0051179), and response to stimulus (GO:0050896). Using the Database for Annovation, Visualization, and Integrated Discovery (DAVID), we identified 85 unique annotation clusters in neonates and 115 clusters in infants that were differentially regulated, as compared to the adolescent/young adult group (**Supplemental Table 6**). Annotation clusters that were overrepresented in both neonates and infants included those associated with the extracellular matrix (e.g., KW-0272, GO:0005201, GO:0031012, GO:0030198, GO:0030020, GO:0005604), collagen (e.g., KW-0176, GO:0005581, GO:0030199), and bioenergetics (e.g., KW-0443, KW-0274, KW-0560, HSA01100). Annotation clusters overrepresented in neonates were predominately related to cell cycle and cell division (e.g., KW-0131, KW-0132, KW-0498, GO:0000070, GO:0072686) (**Figure 2B**), while those overrepresented in infants were related to heart rate regulation (GO:0002027), calcium handling (HSA04020), muscle structure, organization, and contractile function (GO:0030018, KW-0514, HSA05410) (**Figure 2F**).

**Figure 2.**
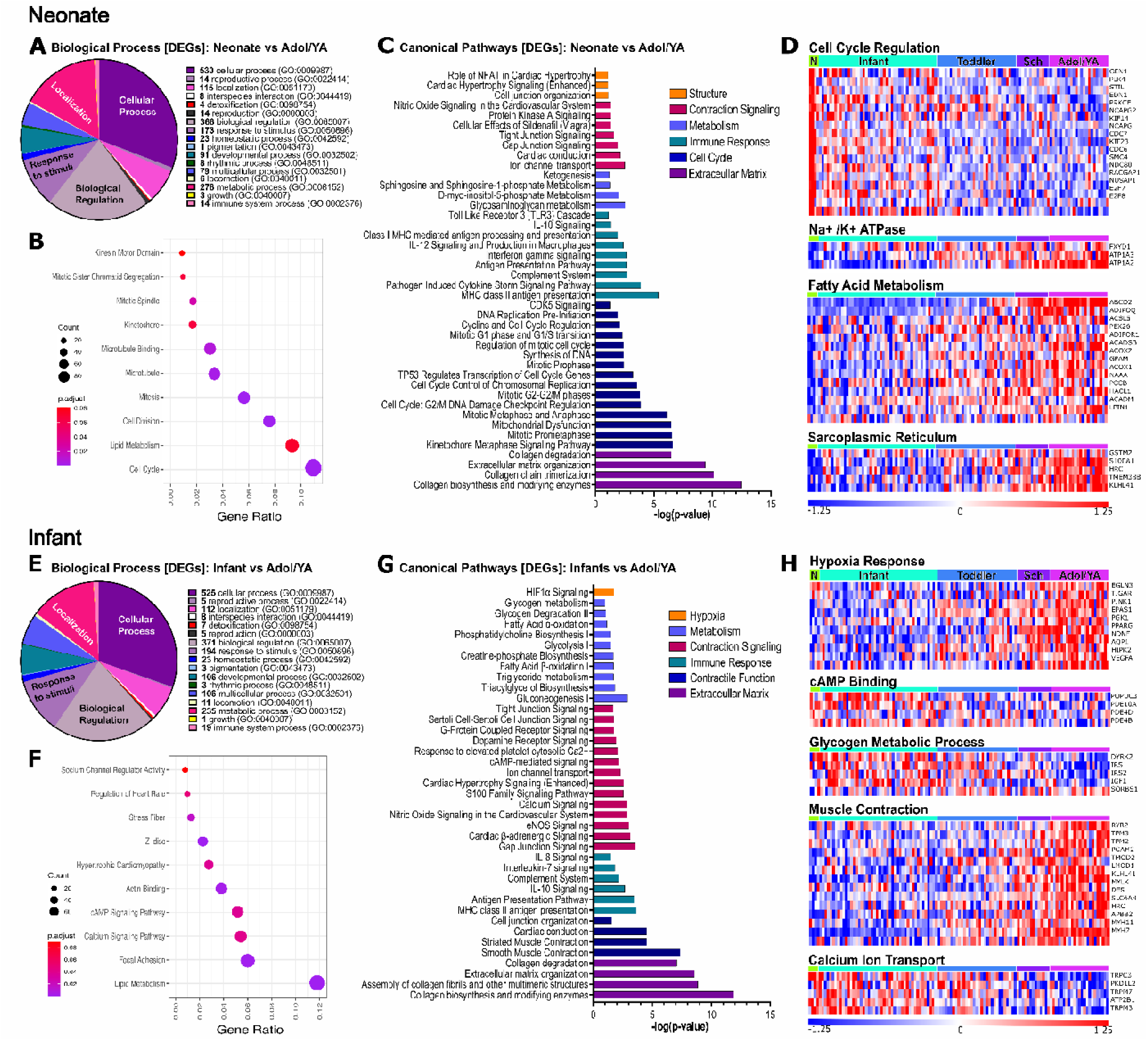
Gene ontology and pathway analysis of differentially expressed genes in neonate group **(Top)** or infant group **(Bottom)**, relative to adolescent/young adults (Adol/YA). **(A, E)** PANTHER analysis of differentially expressed genes into top biological process categories. **(B, F)** Subset of annotation clusters (identified using DAVID) that were overrepresented in neonates or infants, relative to adolescent/young adults. Number of genes per cluster is indicated by the dot size, and Bonferroni-adjusted p-value by the color. **(C,G)** Subset of canonical pathways (identified using Ingenuity Pathway Analysis) overrepresented in neonates or infants, relative to adolescent/young adults. Pathways were filtered using a Fisher’s exact test, with an adjusted p-value (0.1). **(D,H)** Subset of gene ontologies (identified using Enrichr) that are significantly up- or down-regulated in neonates or infants. Heatmap shows each gene (row) for each patient (column) within the specified gene ontology; signal intensity was log_10_ transformed and Z-score normalized on a per gene basis.

Using Ingenuity Pathway Analysis, we identified 252 enriched canonical pathways in neonates and 319 pathways in infants, as compared to the adolescent/young adult group (**Supplemental Table 7**). Canonical pathways that were enriched in both neonates and infants included those associated with contraction signaling (e.g., tight junctions, gap junctions, nitric oxide, ion channel transport), extracellular matrix (e.g., collagen degradation, extracellular matrix organization, collagen biosynthesis), and immune response (e.g., complement system, IL-10 signaling, antigen presentation pathway, MHC class II antigen presentation) (**Figure 2C,G**). Enriched pathways in neonates also included those associated with the cell cycle (e.g., synthesis of DNA, cyclins and cell cycle regulation, regulation of mitotic cell cycle), cell structure (e.g., cell junction organization, cardiac hypertrophy signaling, NFAT in cardiac hypertrophy), and metabolism (e.g., ketogenesis, spingosine metabolism, glycosaminoglycan metabolism). While enriched pathways in infants included hypoxia signaling (e.g., HIF1a signaling), contractile function (e.g., cardiac conduction, striated muscle contraction, cell junction organization), and metabolism (e.g., glycogen metabolism and degradation, fatty acid oxidation, glycolysis). Using the Enrichr analysis tool, we identified 128/273 gene ontologies associated with down/upregulated genes in neonates and 225/39 gene ontologies in infants – as compared to adolescent/young adults (**Supplemental Table 8**). A subset of those gene ontologies is shown in **Figure 2D, H**. Neonates overexpressed genes associated with the cell cycle process and cell proliferation (e.g., *CDC6, CDC7, PLK4*), and underexpressed genes associated with sodium/potassium ATPase activity (e.g., *ATP1A3, ATP1A2, FXYD1*), fatty acid metabolism (e.g., *ACOX1, LPIN1, ADIPOQ*), and the sarcoplasmic reticulum (e.g., *S100A1, SLN, HRC;* **Figure 2D**). In comparison, infants overexpressed genes associated with cAMP signaling (e.g., *PDE4D, PDE4B, PDE10A*), insulin signaling and glycogen metabolism (e.g., *IGF1, DYRK2, IRS1*) and underexpressed genes related to muscle contraction (e.g., *RYR2, MYLK, MYH7, TMP2, TMP3;* **Figure 2H**). Signal intensities for 16 selected genes are shown for each of the five age groups in **Supplemental Figure 2**. As expected, genes related to contractile function and hypoxia-response signaling are upregulated with age – while genes related to cell cycle regulation decreased with age. Notably, although patients were binned into five predetermined age groups, intergroup variability was also evident.

### Genes Encoding Cardiac Ion Channels and Calcium Handling Proteins

Mathematical models of cardiomyocyte electrophysiology and calcium dynamics have recently evolved to evaluate heterogeneities related to sex, age, and/or disease. When ion channel conductances cannot be measured directly, one approach is to assume that conductances are proportional to the gene expression profile of a patient cohort (e.g., male vs female)^25^ or individual patient^26^. Accordingly, we used patient-normalized mRNA expression to generate conductances for the associated ionic currents used in the Grandi model^23^. As such, we refer to these values as they relate to ionic current - while acknowledging the limitation that the relationship between mRNA expression and current conductance does not take into account translation and posttranslational modifications.

Signal intensities for 20 genes involved in cardiac electrophysiology and intracellular calcium handling are shown in **Figure 3A**, with patients binned into each of the five age groups. Age-dependent adaptions in ionic currents followed a variety of trends. A few currents followed a positive linear trend across postnatal development, including *I*_tof_ (*KCND3*), *I*_Kr_ (*KCNH2*), and *I*_KAch_ (*KCNJ3*). While others changed at specific phases of development, including both positive shifts in *I*_RyR_ (*RYR2*) and negative shifts in *I*_K1_ (*KCNJ2/4/12*) and *I*_NCX_ (*SLC8A1*). Our results suggest that *I*_Kr_ remains a major repolarizing current throughout development, as the balance between *I*_Kr_ and other repolarizing currents did not change (*I*_Kr_:*I*_tof_) or even increased with older age (*I*_Kr_:*I*_Kur_ and *I*_Kr_:*I*_Ks_ and *I*_Kr_:*I*_K1_; **Figure 3A**). Almost all gene-current pairs were correlated, and interestingtly, all negative correlations were linked to either *I*_Kur_ or *I*_K1_ (**Supplemental Figure 3**). Other changes between age groups were not considered significant (*I*_Na_, *I*_NaK_, *I*_CaL_, *I*_Ks_, *I*_Kp_, *I*_Kur_, *I*_ClCa_, and *ATP2A2*). Linear trends between age and ionic current expression are illustrated in **Figure 3B**. Expression profiles were partially validated using qRT-PCR for a subset of genes and patient samples (n=24 patients, due to limited tissue resource material). Within this sample subset, we noted congruence between microarray and qRT-PCR results – as *I*_tof_ (*KCND3*) and *RYR2* increased with age, and *I*_NCX_ (*SLC8A1*) had a biphasic profile (**Supplemental Figure 4**). We also evaluated sex-specific differences in gene expression across the entire population (**Figure 4A**) and with separation into five specific age groups (**Figure 4B, Supplemental Figure 5**). We observed increased *I*_Na_ (*SCN5A*) in the entire male population, which may be influenced by a significant increase in *SCN5A* expression in toddler males versus females. We also observed increased SERCA (*ATP2A2*) in the entire male population, which was not attributed to any one specific age group. In age-specific groups, *I*_CaL_ (*CACNA1C*) was increased in school-aged males vs females, while *I_KAch_* (*KCNJ2*) was increased in both toddler and school-aged males – as compared to age-matched females. Sex-specific comparisons were not possible in the neonatal age group due to sample size (n=1 male, n=3 female acyanotic CHD patients).

**Figure 3.**
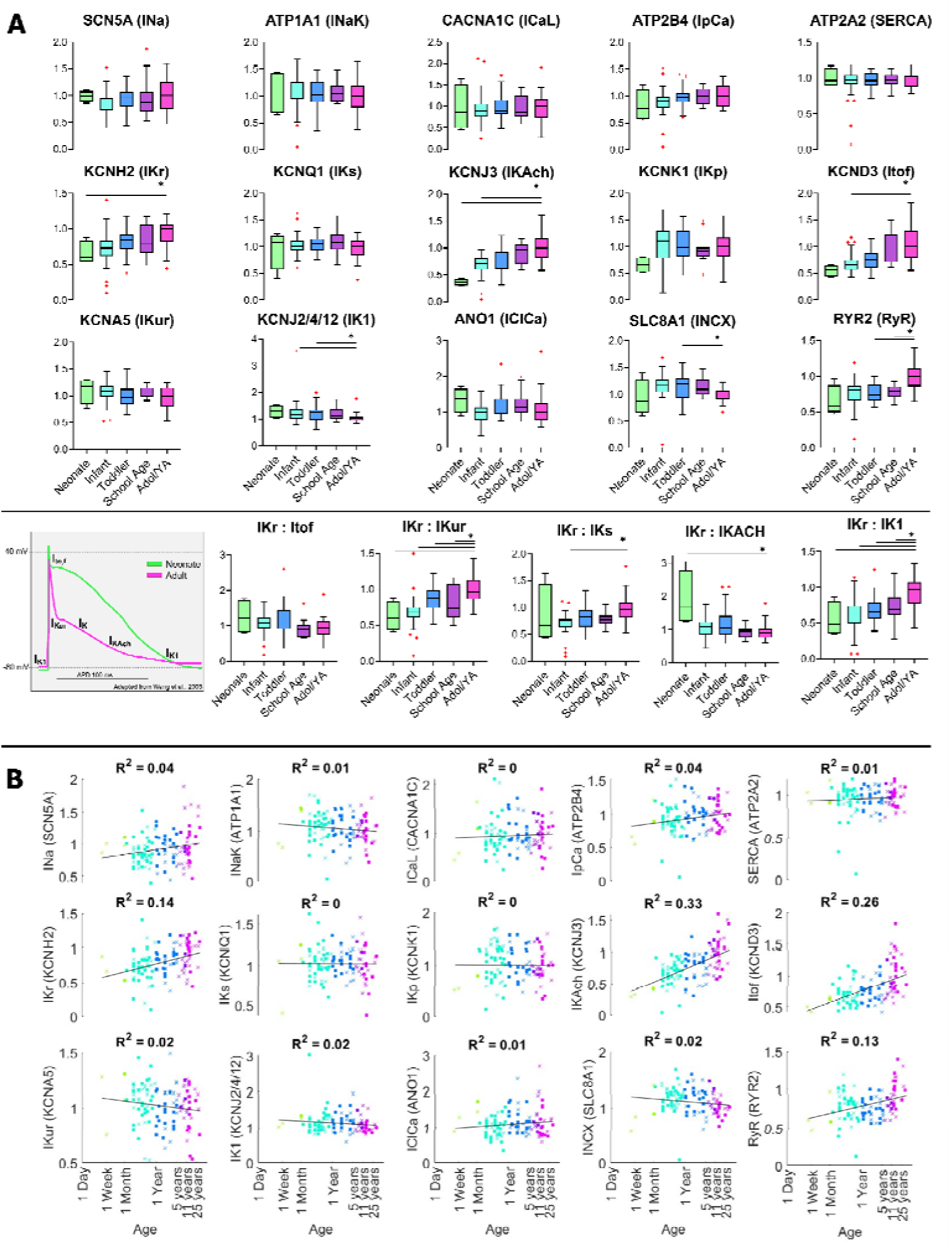
**(A) Top:** Genes related to ionic currents involved in the cardiac action potential and/or intracellular calcium handling. **Bottom:** Experimental action potential recording from neonatal and adult atrial cardiomyocyte (adapted from Wang et al.), with key potassium currents denoted. The relative contribution of each repolarizing current (across each age group), approximated as a ratio to *I*_Kr_ (*KCNH2*). Box plots denote the median expression level (line), interquartile range (box), and outliers (red cross) for each of the five age groups. Note: units are arbitrary, as the raw signals are normalized to the median value of the adolescent/young adults (Adol/YA) group, for each gene. Statistical analysis between each younger age group versus Adol/YA determined by one-way ANOVA with Holm-Sidak test (parametric data) or Kruskal-Wallis with Dunn’s test (non-parametric data); *p<0.05 **(B)** Gene expression data presented continuously, as a function of age (days); x-axis displayed as natural log scale. Results of linear regression analysis shown, with corresponding R^2^ value. Each individual patient is shown (males denoted by “□” and females by “×”).

**Figure 4.**
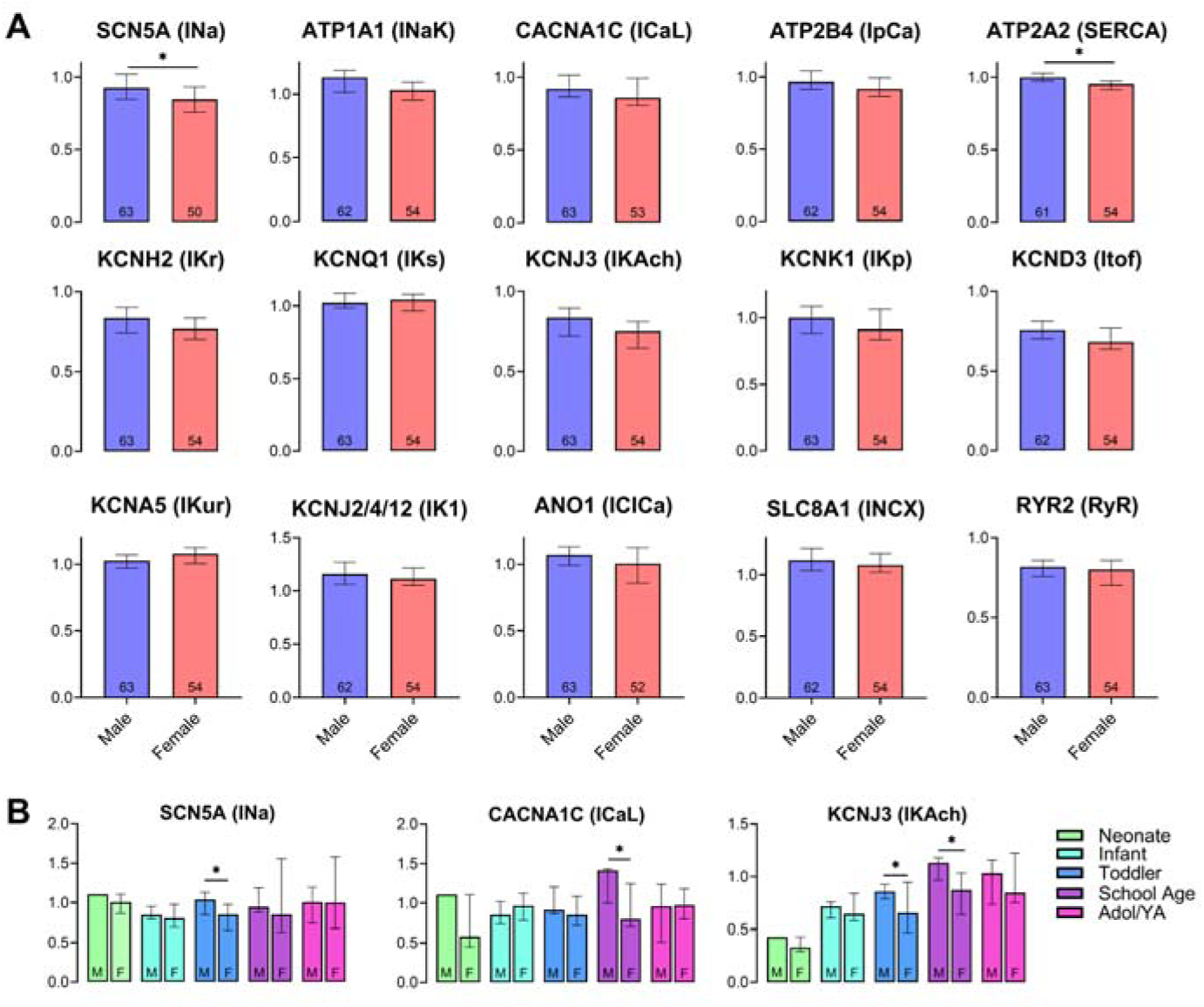
**(A)** Sex-specific differences in gene expression, across the entire population of patients. Note: units are arbitrary, as the raw signals are normalized to the median of the population (n=117). Outliers identified by ROUT method (0.01 false discovery rate); sample size indicated in each bar graph. **(B)** Sex-specific differences in gene expression within each of the five age groups. Note: units are arbitrary, as raw signals are normalized to the median of the oldest age group (adolescent/young adults; Adol/YA). Males denoted with “M” and females with “F” of each pair. All samples are included to maximize sample size. Statistical analysis not feasible for the neonatal group (n=1 male, n=3 female). Data represented as the median ± 95% confidence interval. Statistical analysis by two-tailed Student’s t-test (parametric) or Kolmogorov-Smirnov test (non-parametric), *p<0.05.

### Age-dependent Alterations in Cardiac Electrophysiology and Calcium Handling

Example simulations of patient-specific cell populations are illustrated in **Figure 5A-C**, with one individual patient depicted for each of the five different age groups. Quantitatively, the atrial action potential and calcium transient shapes were maintained across individual patients; however, there were slight and distinct differences in morphology based on the patient-specific conductance scaling factors. Further, there was significant inter-patient variability in the prevalence of “healthy” action potentials versus action potentials with repolarization abnormalities. The range of calcium amplitudes (peak calcium minus diastolic calcium) varied between patients within an age group, but, the majority of simulations were within a range ∼100 nM.

**Figure 5.**
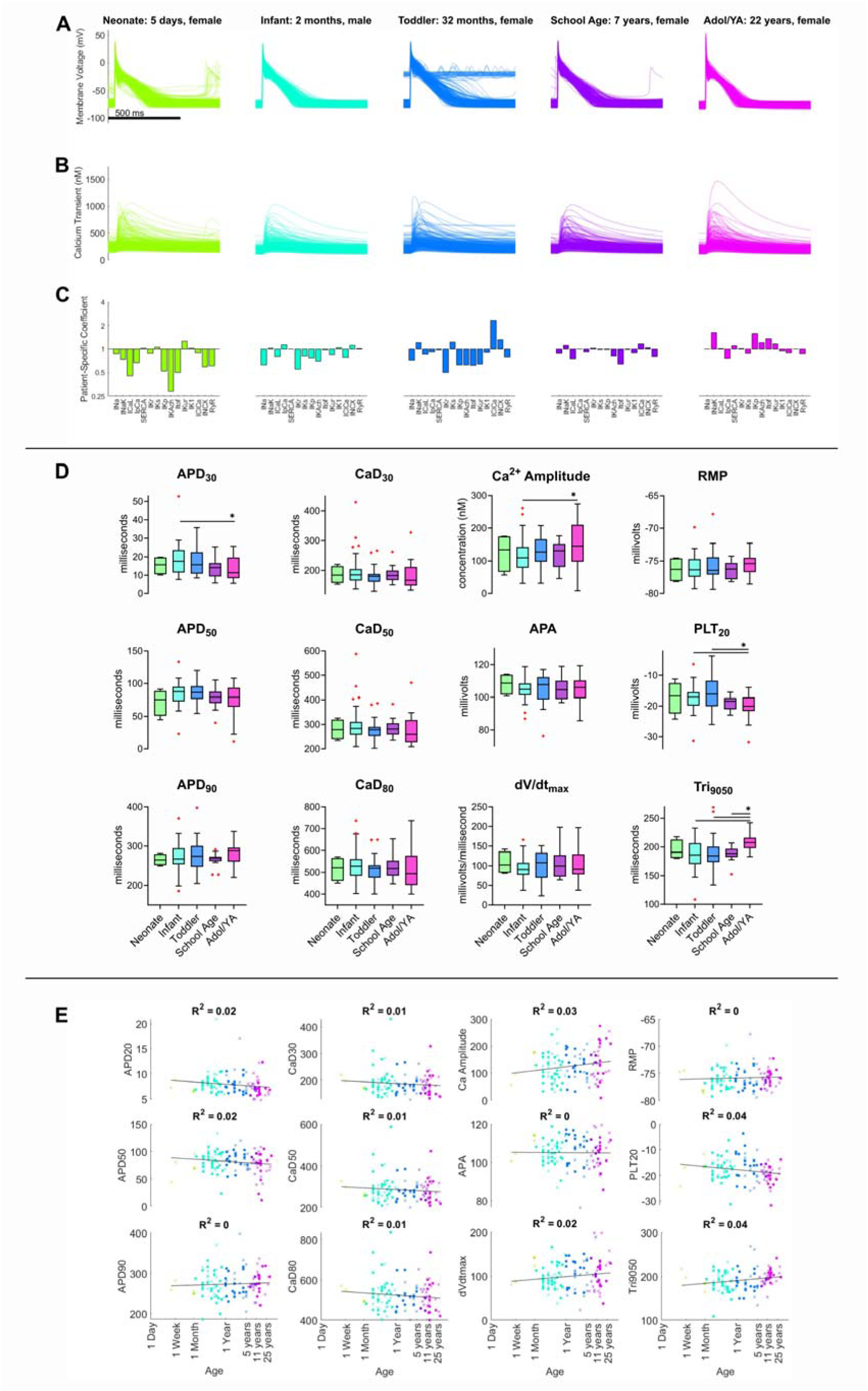
**(A)** Simulated action potentials from a single patient’s population of cells. One individual patient is depicted for each of the five age groups. 500 msec scale bar. **(B)** Simulated calcium transients from a single patient’s population of cells; same scale. **(C)** Simulations are based on patient-specific gene expression (normalized to median value of the adolescent/young adult (Adol/YA) cohort), representing patient-specific current conductance coefficients. **(D)** Age-specific differences in action potential and calcium transient biomarkers, which were calculated based on the maximum values over the last 5 beats for all cells within a patient population (excluding cells with abnormal action potentials). Box plots denote the median expression level (line), interquartile range (box), and outliers (red cross) for each of the five age groups. Statistical analysis between each younger age group versus Adol/YA determined by one-way ANOVA with Holm-Sidak test (parametric data) or Kruskal-Wallis with Dunn’s test (non-parametric data); *p<0.05 **(E)** Action potential and calcium transient biomarkers presented continuously, as a function of age (days); x-axis displayed as natural log scale. Results of linear regression analysis shown, with corresponding R^2^ value. Each individual patient is shown (males denoted by “□” and females by “×”). Abbreviations: action potential duration at 30% (APD_30_), 50% (APD_50_) and 90% repolarization (APD_90_), plateau potential at 20% of APD_90_ (PLT_20_), action potential amplitude (APA), resting membrane potential (RMP), maximum rate of depolarixation (dV/dt_max_), action potential triangulation (Tri_9050_), calcium transient duration at 20% (CaD_20_), 50% (CaD_50_), and 80% (CaD_80_), and the calcium transient amplitude (Ca^2+^ Amplitude).

After completing patient-specific simulations, we quantified action potential and calcium transient biomarkers. Biomarker data for each of the five age groups are illustrated in **Figure 5D**, and linear regression analysis (individual patient age vs biomarker measurement) is shown in **Figure 5E**. Age-group differences in APD_30_ (infant vs adolescent/young adult), Tri_9050_ (infant, toddler, and school age vs adolescent/young adult), PLT_20_ (infant, toddler vs adolescent/young adult), and Ca^2+^ amplitude (infant vs adolescent/young adult) were statistically significant. Our results are in agreement with observations by Wang et al., which noted that the action potential shape is more triangular and the plateau potential is more negative in adult versus pediatric atrial cardiomyocytes^27^. Based on linear regression analysis, the PLT_20_ was primarily determined by *I*_NCX_ (positive correlation, R^2^=0.52), but also linked to *I*_CaL_, *I*_NaK_, and SERCA (positive correlation, R^2^ = 0.34, 0.31, and 0.22 for each current, respectively; **Supplemental Figure 6**). *I*_NCX_ is primarily active during phase 2 of the action potential^28^ thus playing a key role in determining the plateau potential. Similar to the experimental results by Wang et al., we also observed APD_30_ shortening with increasing age, which was driven primarily by *I*_Na_ (negative correlation, R^2^ = 0.27) and *I*_tof_ (R^2^ = 0.24; **Supplemental Figure 7**). In our study, the strongest predictor of action potential triangulation was an age-dependent decrease in *I*_K1_ (R^2^ = 0.56, **Supplemental Figure 8**). This observation was unexpected, as *I*_K1_ was higher in the infant and toddler age groups compared to adolescent/young adults – with a less triangular action potential observed in the younger patient groups. The cause of this relationship may be attributed to the role of *I*_K1_ in action potential termination, as illustrated by a moderate correlation with APD_90_ (R^2^ = 0.43, **Supplemental Figure 9**). Our findings suggest that spontaneous activity in pediatric cardiomyocytes may be attributed to a current other than (or in addition to) *I*_K1_ – such as *I*_NCX_ or *I*_f_, as observed in other immature cardiomyocyte models^29^. And although *I*_tof_ (*KCND3*) increased with age, we did not find a meaningful relationship between *I*_tof_ and APD_90_ (R^2^ = 0.01) or Tri_9050_ (R^2^ = 0.02), suggesting that changes in the transient outward current alone are not sufficient to alter action potential triangulation. Finally, we note that others have observed significantly slower and smaller calcium transients in pediatric cardiomyocytes^12,30^. We observed a statistically significant increase in calcium amplitude with older age (**Figure 5**), which can also be influenced by structural changes to the cardiomyocyte (i.e., sarcoplasmic reticulum calcium content, t-tubule development)^7^ that are not included within this computational model.

We also evaluated sex-specific differences in action potential and calcium transient biomarkers (**Figure 6**). Across the entire population, APD_30_ was shorter and dV/dt_max_ trended higher (p=0.08) in males compared to females (**Figure 6A**). Within specific age groups, APD_30_ trended shorter in male vs female infants (p=0.07), while the school age group presented with sex-specific differences in intracellular calcium handling – with males having a larger calcium transient amplitude and shorter calcium transient duration (CaD_30_, CaD_50_, CaD_80_; **Figure 6B, Supplemental Figure 10**). Sex-specific comparisons were not possible in the neonatal age group due to sample size limitations (n=1 male, n=3 female).

**Figure 6.**
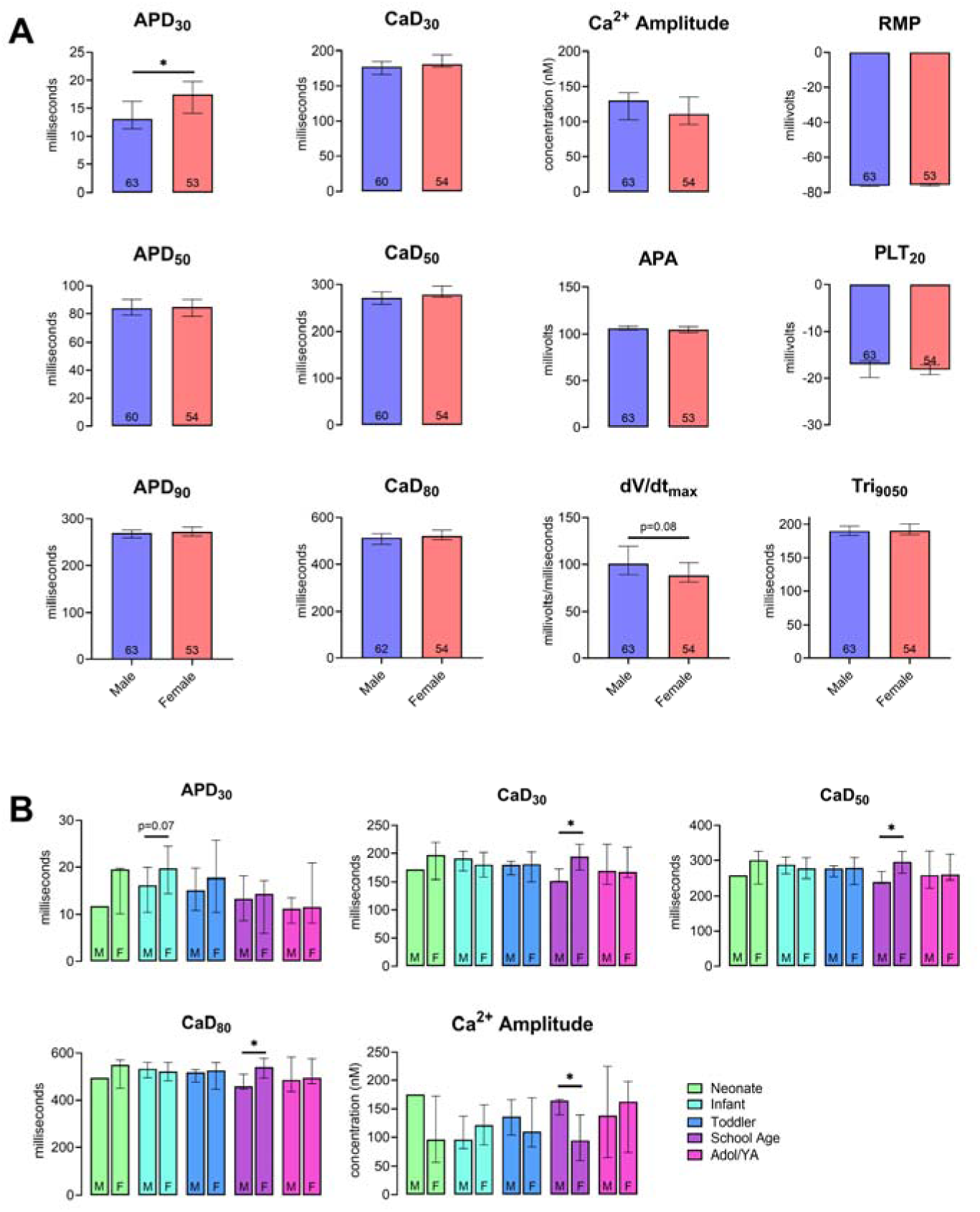
**(A)** Sex-specific differences in action potential and calcium transient biomarkers, across the entire population of patients. Outliers identified by ROUT method (0.01 false discovery rate); sample size indicated in each bar graph. **(B)** Sex-specific differences in action potential and calcium transient biomarkers within each of the five age groups. Males denoted with “M” and females with “F” of each pair. All samples are included to maximize sample size. Statistical analysis not feasible for the neonatal group (n=1 male, n=3 female). Data represented as the median ± 95% confidence interval. Statistical analysis by two-tailed Student’s t-test (parametric) or Kolmogorov-Smirnov test (non-parametric), *p<0.05. Abbreviations: adolescent/young adults (Adol/YA), action potential duration at 30% (APD_30_), 50% (APD_50_) and 90% repolarization (APD_90_), plateau potential at 20% of APD_90_ (PLT_20_), action potential amplitude (APA), resting membrane potential (RMP), maximum rate of depolarixation (dV/dt_max_), action potential triangulation (Tri_9050_), calcium transient duration at 20% (CaD_20_), 50% (CaD_50_), and 80% (CaD_80_), and the calcium transient amplitude (Ca^2+^ Amplitude).

### Simulation Prediction of Patient Arrhythmias

Within the simulated patient populations, the prevalence of abnormal action potentials varied within and between age groups (**Figure 7**). We did not observe an age-specific trend in the arrhythmia score. Next, we investigated the relationship between action potential abnormalities in the simulated patient-specific population with documented clinical arrhythmias. Note that clinical arrhythmia data was only available for a subset of patients: 90 of 117 patients included postoperative arrhythmia data, and of those, 86 of 90 patients had preoperative arrhythmia data. We calculated the receiver operative characteristics (ROC) curve by varying the arrhythmia score threshold to predict the presence or absence of a patient arrhythmia (preoperative and postoperative grouped and ungrouped arrhythmias); the predictive power was evaluated by the area under the curve (AUC). No arrhythmia was predicted with an AUC ≥0.7 (**Supplemental Table 4**). The simulated arrhythmia score provided the best predictive power for preoperative premature atrial contractions (AUC = 0.673); however, there were only two arrhythmias in this group. The second best predictor was for postoperative atrioventricular block (AUC = 0.656, **Figure 7**). Modest predictive power was not entirely unexpected; the simulations used in this study incorporate genetic variability within a patient population – but we acknowledge that this is still an oversimplification relative to patient anatomy and physiology (see Limitations).

**Figure 7.**
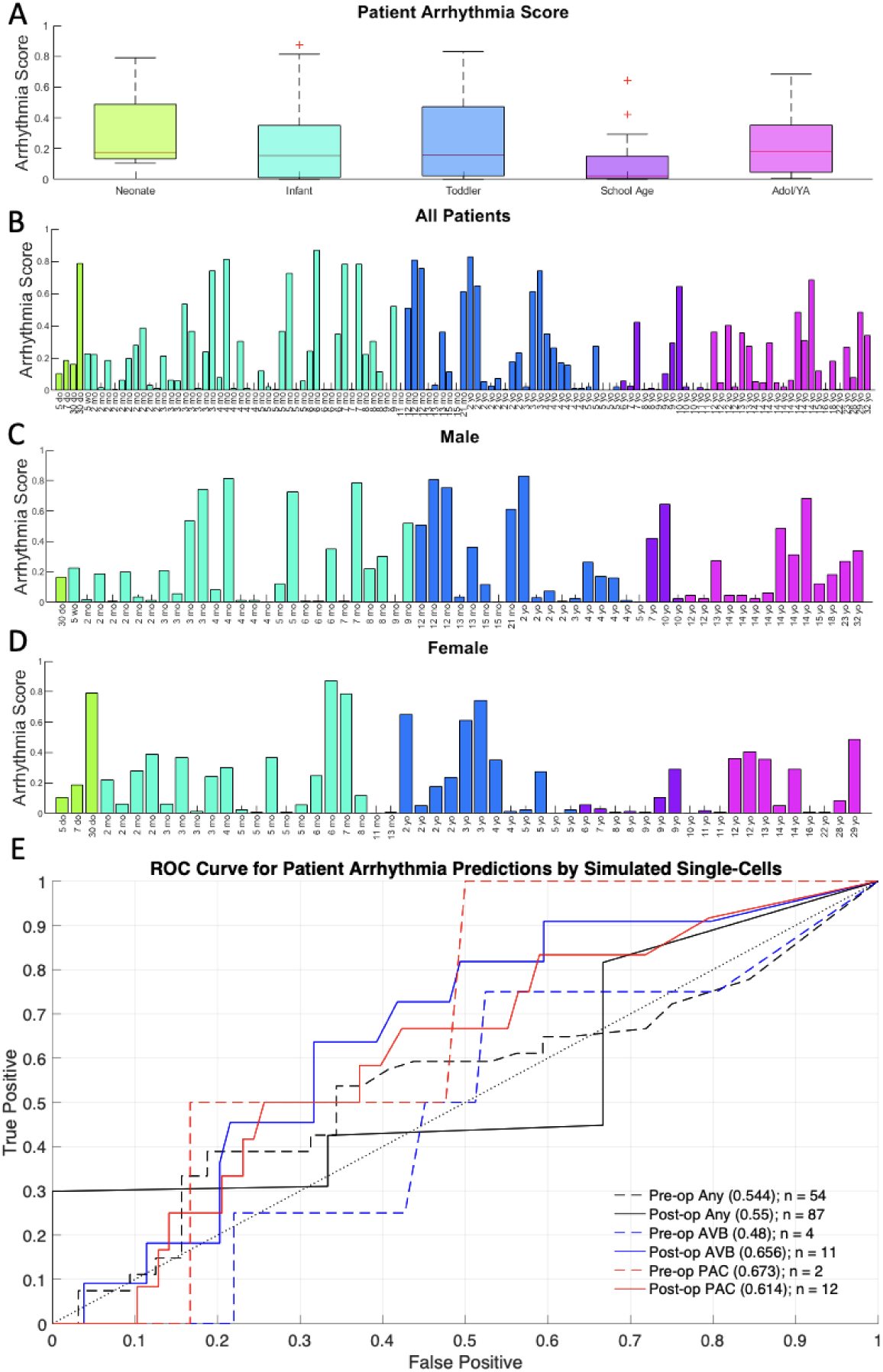
Arrhythmia prediction analysis. **(A)** A normalized single cell “arrhythmia score” was calculated for each patient to quantify irregular electrophysiology, manifesting as abnormal as abnormal action potentials in simulations of specific patients. Arrhythmia predictions were assessed by generating receiver operating characteristic (ROC) curves based on the arrhythmia score threshold, with the area under the ROC curve (AUC) indicating the predictive power of the arrhythmia score for each arrhythmia group or sub-group. All patient arrhythmia scores are presented. **(B-D)** Arrhythmia score for all patient subgroups, organized by age (B) or age- and sex-specific (C,D). The arrhythmia score reflects the fraction of the patient population affected by repolarization abnormality, including no repolarization, early after depolarizations, delayed after depolarizations, or an elevated resting membrane potential. **(E)** ROC curve for predicting patient arrhythmia; ground truth was based on the criteria of any preoperative arrhythmia, any postoperative arrhythmia, preoperative atrioventricular block (AVB), postoperative AVB, preoperative premature atrial contractions (PAC), or postoperative PAC. Postoperative arrhythmias were those recorded within the first five days following surgery. The AUC value is provided in parentheses, followed by the number of patients presenting with the specified arrhythmia. Abbreviation: adolescent/young adults (Adol/YA)

## Discussion

The human heart continues to develop into the first decade of life; yet our current knowledge of human cardiomyocyte maturation and changes in action potential morphology is hindered by the scarcity of healthy heart tissue for biomedical research, especially in the neonatal-pediatric age range. In this study, we reaffirm age is an important biological variable driving changes in global gene expression of human atrial tissue, particularly within the first few years after birth. Age-dependent differences in genes involved in contractile function, hypoxia, and cell cycle regulation were most prominent in patients under 1 year of age compared to patients ≥ 12 years old. Adaptations in some key cardiac ion channel and calcium handling genes were also differentially expressed in our younger patient cohort, which coincides with well-documented developmental changes in electrocardiogram indices^31^. Finally, we translated gene expression into conductances for the associated cardiac ionic currents, where simulations were used to predict developmental and sex-specific changes in action potential and calcium transient morphology.

As cardiomyocytes mature and acquire structural organization, it can become disadvantageous to disassemble (and later reassemble) sarcomeric structures during cell division. Accordingly, cardiomyocyte maturation includes a transition from hyperplasia to hypertrophic growth – but the exact timing in humans is unclear. We demonstrated that neonates and infants overexpress cell-cycle related genes – including *MKI67*, a key biomarker of cell proliferation. This aligns with prior work reporting a decrease in the number of Ki67-positive cardiomyocytes after three months of age – demonstrating a narrow window of proliferation in human atrial cardiomyocytes^8^. Similar trends have been reported in human ventricular tissue, wherein the percentage mitotic cardiomyocytes is ∼11-fold higher in younger samples (<1 year old) and the total number of ventricular cardiomyocytes is generally established within the first few months after birth^32,33^. These findings raise important clinical questions – as the timing of CHD corrective surgery (and the recovery process) could be influenced by a narrow window of cardiac muscle regeneration. Indeed, several surgeons prefer early primary repair (versus palliative options) to maximize on the regenerative potential of the neonatal myocardium, even though this is technically more challenging^34^.

Metabolic remodeling also supports the transition from the intra- to extrauterine environment, as the neonatal heart adjusts to an increasing workload by gradually becoming more dependent on fatty acids as a main energy source. We found that genes associated with fatty acid utilization are overexpressed in older, adolescent/young adult patients. This is in agreement with the study by Yatscoff et al., which reported an age-dependent increase in proteins involved in fatty acid oxidation in ventricular cardiomyocytes^35^. Metabolic remodeling is influenced by oxygen saturation^36^, and as such, postnatal transition to an oxygen-rich environment involves changes in the expression of hypoxia-responsive genes. As one example, we found that neonatal tissue samples overexpress *EPAS1,* the gene encoding hypoxia-inducible factor 2α - a transcription factor that mediates cellular responses to hypoxia. Reduced hypoxia signaling (via HIF1α) has been linked to cardiac metabolic remodeling in both humans and rodents^36^.

The structural organization of developing cardiomyocytes also impacts its contractile efficiency, which is linked to sarcomeric isoform switching (from fetal to adult isoforms). We observed an age-dependent increase in genes associated with contractile kinetics – including myosin heavy chain 7 (*MYH7*) or β-MHC. Cummins & Lambert found β-MHC levels increase in atrial tissue during gestation, but reach maximal expression soon after birth, and thereafter the expression plateaus into adulthood (26% relative to α-MHC expression)^37^. Reiser et al. also found an age-dependent change in MHC isoform expression in fetal and non-failing adult atria, but in this study, the left atria predominately expressed α-MHC^11^. Age-dependent shifts in Troponin I (TnI) have also been previously reported, wherein the slow isoform (ssTnI) in neonates is replaced by the cardiac isoform (cTnI) in adults^9,10,13^. We detected increased *TNNT2* (gene encoding troponin T) expression between neonates and adults, while *TNN1* (gene encoding ssTnI) tended to decrease. We theorize that sarcomeric isoform switching is likely more pronounced in ventricular tissue samples (not examined in this study), due to its more critical role in contraction. With structural maturation, cardiomyocytes also begin to develop protein-rich (e.g., NCX, L-type calcium channels) transverse tubules that facilitate dyadic coupling with the sarcoplasmic reticulum^38^. We report that *SLC8A1* (gene encoding NCX) expression appears biphasic, with elevated expression in the intermediate age groups (infant – school age) and reduced expression at the two extreme age groups (neonates and adolescent/young adults). A similar biphasic trend has previously been reported for NCX protein expression^39^. Finally, we observed that ryanodine receptor expression (*RYR2*) is consistent throughout early development (neonatal to school-age) and then increases at older ages (adolescent/young adult). Reduced RyR expression is linked to immature calcium handling, including a slower calcium transient upstroke time^38^.

Cardiac electrophysiology is also modified throughout human development, including an age-dependent change in *I*_tof_^15–18^. To date, a few studies have suggested that *I*_tof_ density is increased in adult versus pediatric cardiomyocytes^16–18,40^ – although others have observed no age-related change^41^. We observed a continually increase in *KCND3* (gene associated with *I*_tof_) from infancy to adolescent/young adulthood. We cannot comment on activation or inactivation properties, but, we can confirm that detectable levels of *KCND3* were found in all tested patient samples. Of interest, both Mansourati & Le Grand^16^ and Crumb et al.^18^ reported age-dependent differences in *I*_tof_, but noted that it was indetectable in some pediatric cardiomyocytes. We also observed an upward trend in *KCNJ3* (gene associated with *I*_KAch_), an acetylcholine-activated potassium current, from neonates through adolescence/young adulthood. To our knowledge, no studies have measured age-dependent changes in this current, but a neurotransmitter switch from catecholamine to acetylcholine does occur during development. The higher heart rate observed in children is also consistent with decreased parasympathetic activity.

In this study, we also report a significant decrease in *KCNJ2/4/12* (genes associated with *I*_K1_) and a significant increase in *KCNH2* (gene associated with *I*_Kr_) with older age. Previous studies have investigated age dependence of *I*_K1_ in both the ventricle^42^ and atria^18^, with conflicting results. Schaffer et al. reported 50% less *I*_K1_ density than previously reported in adults, suggesting that current density increases with age^42^. In contrast, Crumb et al. found that *I*_K1_ density was comparable between pediatric and adult atrial cells^18^. Age-dependent changes in *I*_Kr_ are also understudied in the context of human cardiomyocyte maturation, but developmental studies have been conducted using ventricular tissues from animal models. Using dofetilide to block *I*_Kr_, Wang et al. reported that *I*_Kr_ contribution to the mouse AP is higher in fetal and neonatal tissue and lower in adults^43^. In agreement, Obreztchikova et al. reported that *I*_Kr_ was the dominant repolarizing current in pediatric canine cells^44^. Both conclude that the major repolarizing current moves away from *I*_Kr_ throughout development; however, we found that the balance between *I*_Kr_ and other major repolarizing currents either does not significantly change (*I*_Kr_:*I*_tof_) or increases (*I*_Kr_:*I*_K1_ and *I*_Kr_:*I*_Kur_ and *I*_Kr_:*I*_Ks_) with age in human atrial samples, suggesting that these changes could be species and/or chamber specific. Lower *I*_Kr_ in younger patients could be related to the para/sympathetic activity during development. Younger patients have a higher heart rate and underdeveloped parasympathetic signaling, which could influence their responsiveness to beta-adrenergic activation – a process that increases repolarization from *I*_Ks_ and elevates *I*_K1_^45^. As such, the demand for total repolarization may be predominately provided by *I*_Ks_ in pediatrics, instead of *I*_Kr_ as in adults^46^. Notably, increasing *I*_Kr_ and decreasing *I*_K1_ are both positively correlated with *I*_KAch_ – suggesting that these developmental changes could be linked to parasympathetic innervation.

Finally, in our study we observed only modest sex-specific differences in ion channel expression, action potential metrics, or calcium handling biomarkers. Males had elevated expression of *SCN5A* (gene associated with *I_Na_*), *ATP2A2* (gene encoding SERCA), and *KCNJ3* (gene encoding *I*_KAch_) – which was largely attributed to increased expression in the male toddler and school-age cohort, compared with age-matched females. Within the entire population, males had a shorter APD_30_ and a more positive max rate of depolarization compared to females. Only patients within the school-age group presented with sex-based differences in calcium handling, with a larger calcium amplitude and shorter calcium transient duration measured in males. These results are in agreement with experiments in small mammals, in which sex hormones cause females to develop smaller/slower calcium transients and contractions compared to males ^47–49^. Notably, we did not observe any significant changes in action potential or calcium biomarkers in the mature adolescent/young adult group.

Gaps in our understanding of pediatric electrophysiology can have unintended consequences, particularly when pharmacological therapies are designed and tested using only adult cardiac models: targeting ion channels that may be underdeveloped in the immature myocardium. Further, measurements of action potential and calcium transients require the use of live cardiomyocytes, and as a result, these studies are extremely limited in human cardiomyocytes. Our simulations predict quantitative patient-specific ion channel conductances, estimating atrial action potential and calcium transient shapes across individuals. We report an age-dependent decrease in the AP plateau potential and increase in AP triangulation, which could be associated arrhythmic events. These findings are in agreement with Wang et al., in which adult human atrial cardiomyocytes exhibited a more negative plateau phase and a trend toward shorter action potential duration as compared to pediatric cardiomyocytes (see experimental action potential in **Figure 3A**)^40^.

## Limitations

Our study, while contributing valuable insights, encompasses experimental constraints due to the exclusive use of atrial tissue for our analyses. Recognizing the documented chamber-specific differences in cardiac physiology, our findings may not be readily extrapolated to other cardiac chambers. When possible, future research should explore variations in gene expression and ionic current characteristics across different cardiac compartments in humans. Another potential limitation of our study is our patient cohort, while characterized by acyanotic status, presents with congenital heart diseases. Considerably, our samples do not inherently represent a healthy population, and future studies should aim to compare samples from individuals without congenital heart diseases to enhance the generalizability of our findings. Additionally, our approach involves making assumptions from gene expression to ionic current conductance without an intermediate step involving protein expression. Future investigations should explore incorporating protein-level data to refine the accuracy of the model and provide a more comprehensive understanding of the gene-to-current relationship. Furthermore, we acknowledge that our discussion of the relationship between RNA expression and current conductance does not account for post-transcriptional regulation. Future research should explore the role of post-transcriptional regulation in shaping the ionic current landscape to refine our understanding of the observed relationships. Further, when correlating simulation arrhythmia risk with patient arrhythmia occurrences, poor predictive power was not fully unexpected, as the models used within this study are oversimplified (e.g., lacking cell-cell coupling, anatomical consideration, and electrophysiology of non-atrial cell types). Additionally, evidence indicates that changes in sympathetic and parasympathetic activity (e.g. β-stimulation, *I*_KAch_) play a critical role in changing repolarization reserve throughout development, and should be investigated in future work. Despite initial negative results, the methods used within this study have the potential to increase the accuracy of *in-silico* patient models. Future work incorporating patient-specific electrophysiology into detailed biophysical models can be coupled with virtual whole-heart models created from detailed clinical scans as described by Sung et al. to create patient-specific, anatomically and electrophysiologically accurate, whole-heart models^50^. If successful, this would improve upon current *in-silico* standards of arrhythmia prediction, ablation location optimization, and patient drug response.

## Acknowledgements

We gratefully acknowledge Susan Knoblach, Karuna Panchapakesan, and the Children’s National Research Institute Genomics and Bioinformatics Core for assistance with microarray experiments. We also acknowledge additional members of the cardiac surgery operating room team, including Aybala Tongut, Louise Wilson, Lula Curry, Hyung Mi (Grace) Yang, Sandra Sunderland, Evelyn Ravizee, Delia Angeles, Jeanette Callos, and Blezzy Bote for their assistance with tissue sample procurement.

## Author Contributions

DG, PS, CY, MD, YD, AB screened acyanotic CHD patients for enrollment and preserved atrial tissue samples during cardiac surgery for this biomedical research study. CY, PS, DG, and NGP conceived and designed the human subjects study. SS, DG, JM, NM, SHW, NGP performed experiments, analyzed and interpreted data. SS, JM, DG prepared figures and tables. SS, JM, SHW, and NGP drafted the manuscript. All authors revised the manuscript or provided critical intellectual content, and all authors approved the final manuscript.

## Data Availability

Derived data supporting the findings of this study are available from the corresponding author (NGP) upon request.

