## Supplemental Methods, Figures, Tables 1-5 for "Connecting Transcriptomics with Computational Modeling to Reveal Developmental Adaptations in the Human Pediatric Myocardium"

Human Subjects: Tissue collection and experimental studies were performed in accordance with a Children’s National Hospital Institutional Review Board-approved protocol (IRB#Pro00012146). This minimal risk study involved the collection and preservation of right atrial tissue samples, predominately the right atrial appendage (as medical waste), from patients who underwent open heart surgery at Children’s National Hospital (**Supplemental Table 1,** n=117). Prior to surgery, the clinical characteristics of each patient were assessed by one of the study investigators. Patients with acyanotic CHD (>94% O_2_ saturation) were enrolled in this study if their procedure included removal of right atrial tissue during surgical repair. Donor information included age, gestational age, sex, weight, STAT category, oxygen saturation level, and incidence of pre- or postoperative arrhythmia. Tissue samples were designated to one of five age groups: neonate (0-30 days, n=4), infant (31-364 days; n=46), toddler to preschool (1-5 years, n=31), school age (6-11 years, n=13), and adolescent to young adults (12-32 years, n=23). The gestational age of each infant patient was verified, and in one case a premature infant was included in the neonate group due to age-correction.

Gene Expression Analysis: Cardiac tissue samples were submerged in RNAlater stabilization solution (Invitrogen, Waltham MA USA) and stored at 4°C for <7 days. RNAlater was subsequently removed and samples were then stored at -80°C until RNA extraction. Total RNA was isolated from right atrial tissue (5-30 mg) and processed using a RNeasy fibrous tissue kit with on-column DNase treatment (Qiagen, Germantown MD USA). RNA concentration was determined using a NanoDrop spectrophotometer (Thermo Fisher, Waltham MA USA) and RNA quality was assessed using a Bioanalyzer 2100 (Agilent Technologies, Santa Clara CA USA). Only samples of sufficient quality were used in subsequent microarray experiments (Mean RNA integrity number: 8.6±0.7, concentration: 244±137 ng/μl, RNA ratio: 2.0±0.1). 250 ng of total RNA input was primed for the entire length of RNA, including both poly(A) and non-poly(A) mRNA and reverse transcribed to generate sense-stranded targets that were biotin-labeled using a GeneChip WT Plus Reagent kit, and then hybridized to Human Clariom Arrays (Applied Biosystems, Waltham MA USA) for 16 hours at 45°C. After removing the hybridization cocktail, each array was washed and stained on the Fluidics Station F450, and then scanned using an Affymetrix GeneChip Scanner (GCS 3000 7G; Thermo Fisher). Initial quality control data was evaluated using Affymetrix Transcriptome Analysis Console Software. Microarray data were imported and analyzed using the Transcriptome Analysis Console (Applied Biosystems). To identify differentially expressed genes (DEGs), data sets were compared between age groups using one-way ANOVA with a p-value threshold ≤0.05, fold change cutoff of ≥|1.25|, and a false discovery rate of ≤0.1. For each DEG, the raw signal intensity data underwent log_10_ transformation and Z-score normalization (Z score = (signal intensity – mean signal intensity)/standard deviation of signal intensity) on a per-gene basis. Z-scores were used to display DEGs as a heatmap using TIGR MultiExperiment Viewer^1^. Enrichment analysis was performed on a single rank-ordered gene list via analysis tools, including David Bioinformatics, Panther, Ingenuity Pathway Analysis (Qiagen), and Enrichr^2–8^. Datasets are available via the Gene Expression Omnibus.

Real-Time Quantitative Reverse Transcription PCR (qRT-PCR): Microarray data trends were validated by qRT-PCR, using a limited number of genes and tissue samples. Briefly, RNA was reverse transcribed using a SuperScript VILO cDNA synthesis kit, according to the manufacturer’s instructions (Invitrogen). TaqMan gene expression assays were used for fluorescence-based qRT-PCR via a QuantStudio 7 platform (Applied Biosystems). Gene expression was normalized to the reference genes glyceraldehyde-3-phosphate dehydrogenase (*GAPDH*) and RNA polymerase II polypeptide A (*POLR2A*) using the comparative C_T_ method (or 2^-ΔΔCT^)^9^. Fold change values are reported (average of three technical replicates), with each assay including a minimum of three individual biological replicates.

Computational Model Selection and Approach: Atrial cardiomyocyte electrophysiology was simulated by adapting the Grandi, et al. model^10,11^, motivated by the results of Muszkiewicz et al.^12^ that demonstrated that this model can reproduce the action potential and ionic currents of human atrial cells from the right atrial appendage. To develop patient-specific simulations, we scaled the ionic currents in the Grandi, et al. atrial cell model based on the current-specific gene expression for each individual patient, relative to the median expression level in the adolescent/young adult age group. Given the uncertainty and variability in this older age group, we first developed a generic adult “control” population, unbiased by patient gene data, fit to experimental action potential biomarkers for the right atrial appendage^13^. Each cell in the control population was then adjusted by the patient-specific ionic current scaling factors (based on gene expression level) to generate a population of atrial cells for each individual patient. For each patient, the action potential characteristics of the cell population were then used to quantify patient-specific and age-dependent measurements.

Simulation: Atrial cardiomyocytes were simulated using a stimulation current of 12.5 μA/μF for a duration of 5 ms. Ordinary differential equations for membrane potential and state variables (fully provided in ref. ^10^) were integrated using the build-in MATLAB function ODE15s (MathWorks, Natick MA USA). To mimic a normal adult heart rate, a pacing frequency of 1 Hz (60 beats per minute) for 50 beats was used for all simulations. Although the average human heart rate gradually decreases from infancy to adulthood^14^, in this study, we used a fixed pacing rate to avoid introducing frequency-dependency in the simulated results.

Generation of an Adult Control Population: Prior to developing age-specific models, we first generated a “control” population to represent mature atrial cardiomyocytes by varying the ionic current conductances by a scaling factor (*θ*), as previously performed by our lab^15^ and others^16–18^. This control population represents the variability present in the overall adolescent/young adult population of atrial myocytes. To generate the control population, the maximal conductance of 19 ionic currents and exchangers was varied between 1/4-4x the model baseline using Latin hypercube sampling. This approach generated 30,000 unique single-cell parameter sets within the sampling space; scaling was distributed uniformly on a logarithmic scale, such that there is an equal likelihood an ionic current is scaled to 1/4x (θ = 0.25) or 4x the model baseline (θ = 4). Action potential characteristics were measured for each atrial myocyte (i.e., each parameter set), and only myocytes with characteristics fit to published experimental biomarker values were included in the control population (**Supplemental Table 2**) with the fit defined as two standard deviations from the biomarker mean. Notably, two datasets with distinct action potential duration (APD) values were found in the literature, including Ravens et al. (APD_90_ = 317.41±9.33 ms) and Muszkiewicz et al. (APD_90_ = 107.8±21.9 ms)^12,13^. Since it was not possible to simultaneously fit both experimental datasets, we utilized the Ravens et al. dataset due to its larger sample size (238 vs 29 measurements) to create the control population. In addition, we restricted atrial myocytes included in the control population to those with a calcium transient amplitude >50 nM based on values reported in healthy tissue by Heijman et al.^19^. Collectively, after fitting to all atrial biomarkers, our control population included 240 of 30,000 atrial cells (see **Supplemental Figure 1** for parameter distributions). Deviations from the sampled distribution include higher *θCaL* and *θto,f,* and a lower *θNa.* Importantly, using a minimum amplitude to restrict the calcium transient, the distribution of the calcium transient amplitude closely resembled previously reported experimental values in human atrial cells with 208.7±162.4 nM *in-silico* vs. experimental measures of 183.9±89.9 nM^20^ or 278.1±178.4^21^.

Adaptation of Gene Data for *In-silico* Modeling: Gene selection was based on previously published associations with ionic currents included in the updated Grandi model^10,11^ (**Supplemental Table 3**). Gene-current pairs were primarily taken from a published study in ventricular cells by Smirnov et al.^22^, except as noted below. For each gene and for each patient, the expression level was normalized to the median expression level of the adolescent/young adult patient cohort. The normalized values were subsequently used as coefficients to scale ionic current conductance in each patient-specific simulation. Ionic currents in the model were scaled relative to the adolescent/young adult population, since currents in the Grandi atrial myocyte model were formulated based on adult donor tissue samples^23–28^ and experimental biomarkers that were used to restrict the control population were recorded using adult samples^12^. Since *I*_K1_ is carried by three distinct Kir isoforms, Kir2.1, Kir2.2, and Kir2.3, encoded by *KCNJ2*, *KCNJ12*, and *KCNJ4*, respectively, the averaged expression of the three isoforms was used to scale *I*_K1_^29^. Since atrial myocytes lack the slow transient inward current, the gene encoding the fast component was used (*KCND3*). The remaining genes encoding atrial-specific currents were defined using the *GenBank* database^30^. Genes associated with background currents (representing unknown current sources) were unavailable in the selected patient dataset, and as such, were not included.

Patient-Specific Simulations: A population of cells was generated for each patient by multiplying the patient-normalized gene values by each control cell’s set of scaled conductances in an element-wise fashion. For background currents lacking an associated gene (*I*_Na,b_, *I*_Ca,b_, and *I*_Cl,b_), the control cell conductances were left unchanged. This resulted in a patient-specific set of scaled ionic current conductances for each associated cell in the control population (i.e., all 240 cells), which are simulated to generate a patient-specific population of cells equal in size to the control population. To illustrate this process, if an individual were to have gene expression equal to the median of the adolescent/young adults for all currents, it would result in a cell population identical to the control population. We represented each individual patient with a population of cells, since a single cell (and parameter set) cannot properly recapitulate the electrophysiology of an individual with a high level of certainty. The described population approach accounts for a margin of error for each patient’s predicted electrical activity, and subsequently, to the predicted action potential and calcium transient measurements averaged across each developmental stage.

Biomarker Calculations: Biomarkers were calculated for all cells within each patient population as the maximum value over the last five beats. Biomarkers were selected based on those used to calibrate the control population (**Supplemental Table 2**), as adopted from the referenced experimental study^13^. Collectively, we measured the action potential duration at 30% (APD_30_), 50% (APD_50_) and 90% repolarization (APD_90_), the plateau potential at 20% of APD_90_ (PLT_20_), action potential amplitude (APA), resting membrane potential (RMP), and the maximum rate of depolarization (dV/dt_max_). APD_30_ was used in lieu of APD_20_, in order to provide a direct comparison with experimental data^31^. We also included action potential triangulation (Tri_9050_), which was calculated as the APD_90_ minus APD_50_^12.^ Additionally, we measured biomarkers to characterize intracellular calcium handling, including the calcium transient duration at 20% (CaD_20_), 50% (CaD_50_), and 80% (CaD_80_), and the calcium transient amplitude (Ca^2+^ Amplitude). The median value of each biomarker was calculated for each patient-specific population, excluding cells with an abnormal action potential (described below).

Arrhythmia Risk Assessment and Predictions: The goal of the *in-silico* studies was to predict changes in action potential and calcium transient morphology throughout development. Additionally, it was noteworthy that simulations for specific patients were particularly prone to irregular electrophysiology, with abnormal action potentials defined by failure/no repolarization, early after depolarizations, delayed after depolarizations, or an elevated resting membrane potential (greater than -60mV). To examine this further, we calculated a normalized single-cell “arrhythmia score” for each individual patient, which was defined as the fraction of the cell population with abnormal action potentials. Such an approach was previously employed by Passini, et al. in virtual ventricular cells^17^. Next, we investigated concordance between the prevalence of abnormal action potentials in our *in-silico* studies with the documented arrhythmia incidence for each patient. The latter was performed using a double-blind approach, wherein the predicted single-cell arrhythmia score calculations and the patient arrhythmia history were generated independently. Arrhythmias were assessed preoperatively via electrocardiogram reports, and postoperatively (within first 5 days) via electrocardiogram reports and daily progress or event notes. Patient arrhythmia history was used to define binary classifications for each arrhythmia group. The predicted arrhythmia score values were utilized to generate a receiver operating characteristic (ROC) curve, with points on the ROC curve based on the arrhythmia score threshold. The predictive power of the arrhythmia score was evaluated based on the area under the ROC curve (AUC) for a given arrhythmia group.

Statistical Analysis: The number of biological replicates, statistical test implemented, and type of graphical data depiction (e.g., median with 95% confidence interval) are specified in each figure legend. Data sets were tested for normality using the Shapiro-Wilk test and variance using an F-test (two groups) or Barlett’s test (three or more groups). Normally distributed data were analyzed via one-way ANOVA with/without Welch’s correction (unequal variance), and nonparametric data were analyzed via Kruskal-Wallis ANOVA. For sex-specific analysis, normally distributed data were analyzed via two-tailed t-test with/without Welch’s correction (unequal variance) and nonparametric data were analyzed via Kolmogorov-Smirov test. Linear relationships between age and gene expression were characterized by linear regression analysis, with results illustrated by the coefficient of determination (R^2^). Holm-Sidak or Dunn’s test was used to adjust the significance level for multiple comparisons^32^. Statistical significance is denoted in each figure with an asterisk to signify *p<0.05.

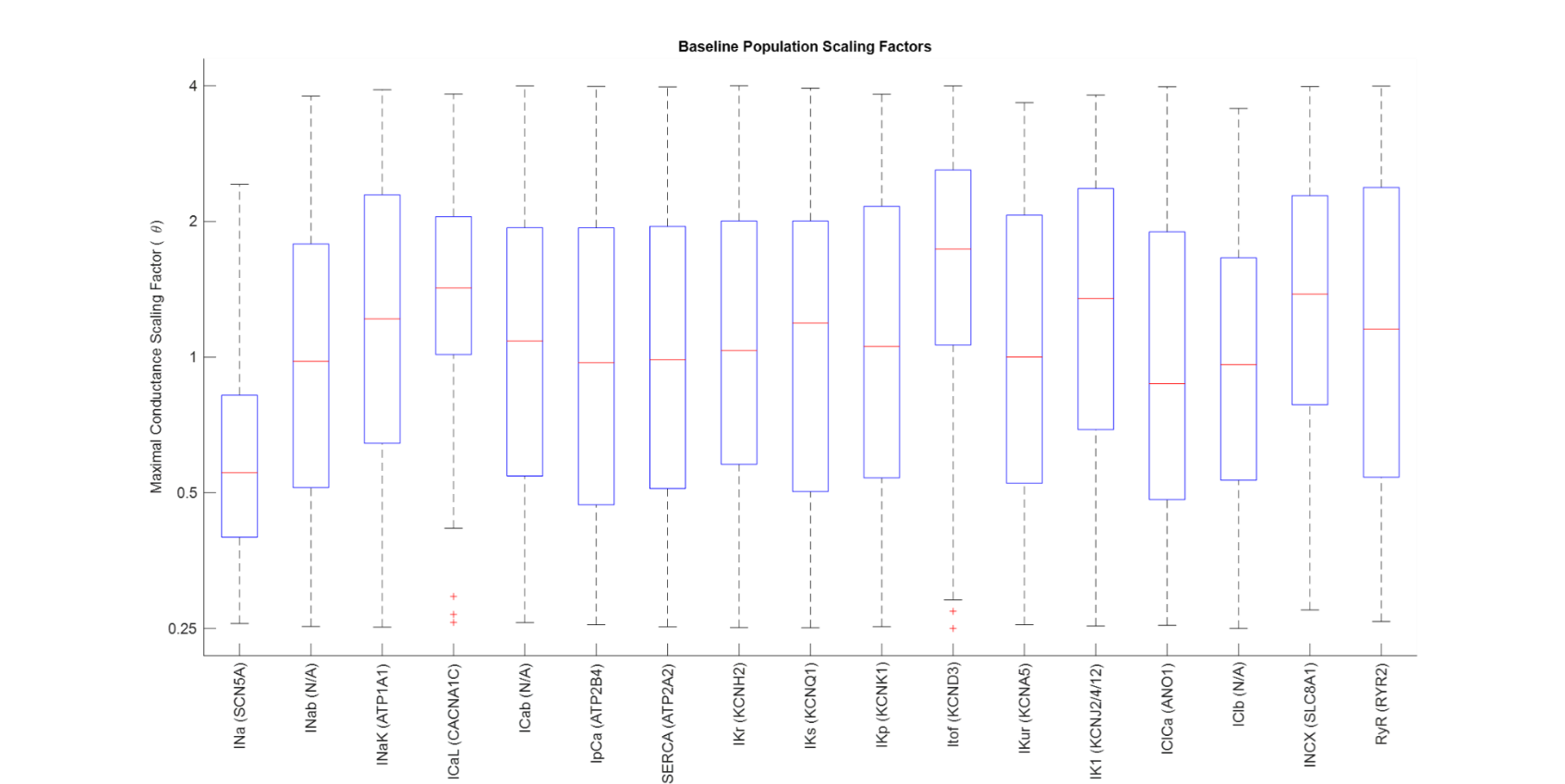

**Supplemental Figure 1.** Distribution of scaling factors in control cells generated using the updated Grandi, et al. model of a human atrial cell, following fit to experimental biomarkers (n = 240 cells). Results indicate a trend towards decreased *I*_Na_, and increased *I*_CaL_, *I*_tof_, *I*_K1_, and I_NCX_ compared to the model baseline. Data shown as the median (red line), interquartile range (blue box), statistical outliers (red crosses).

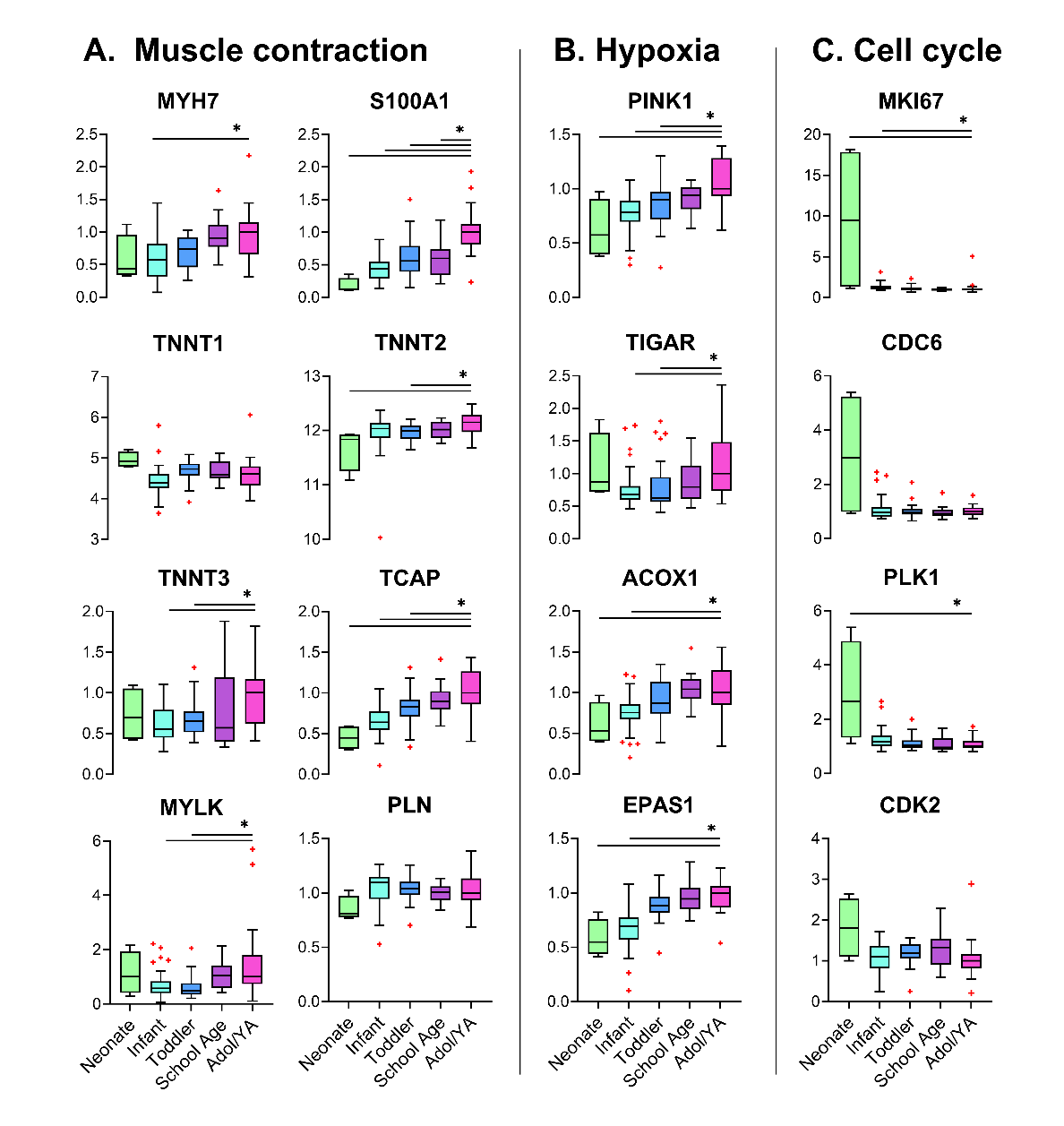
**Supplemental Figure 2.**  Example of three gene ontologies **(A)** muscle contraction, **(B)** hypoxia response, and **(C)** cell cycle that are modified in the neonate and/or infant cohort – as compared to the adolescent/young adult group (Adol/YA). A subset of genes within each gene ontology are shown, denoted by the median expression level (line), interquartile range (box), and outliers (red plus) for each of the five age groups. Note: units are arbitrary, as the raw signals are normalized to the median value of the Adol/YA group, for each gene. Statistical analysis between each younger age group versus Adol/YA determined by one-way ANOVA with Holm-Sidak test (parametric data) or Kruskal-Wallis with Dunn’s test (non-parametric data). *p<0.05

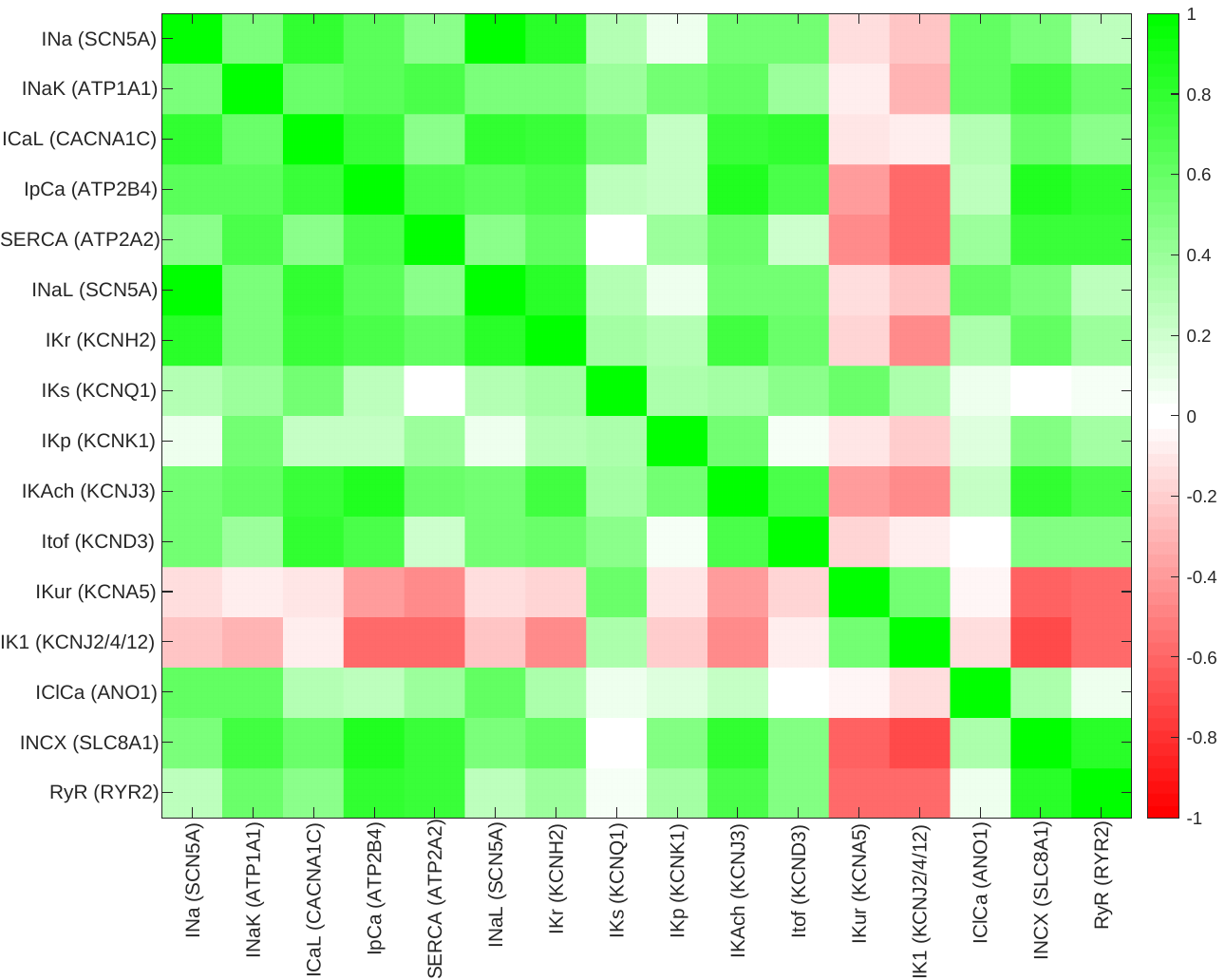

**Supplemental Figure 3.** Pearson’s correlation coefficient between the normalized mRNA expression illustrated in Figure 3, across the entire patient population. We note that changes in repolarization reserve (*I*_Kur_, *I*_K1_) are associated with the development of parasympathetic signaling in the heart, as illustrated by *I*_KAch_, an acetylcholine-activated potassium current.

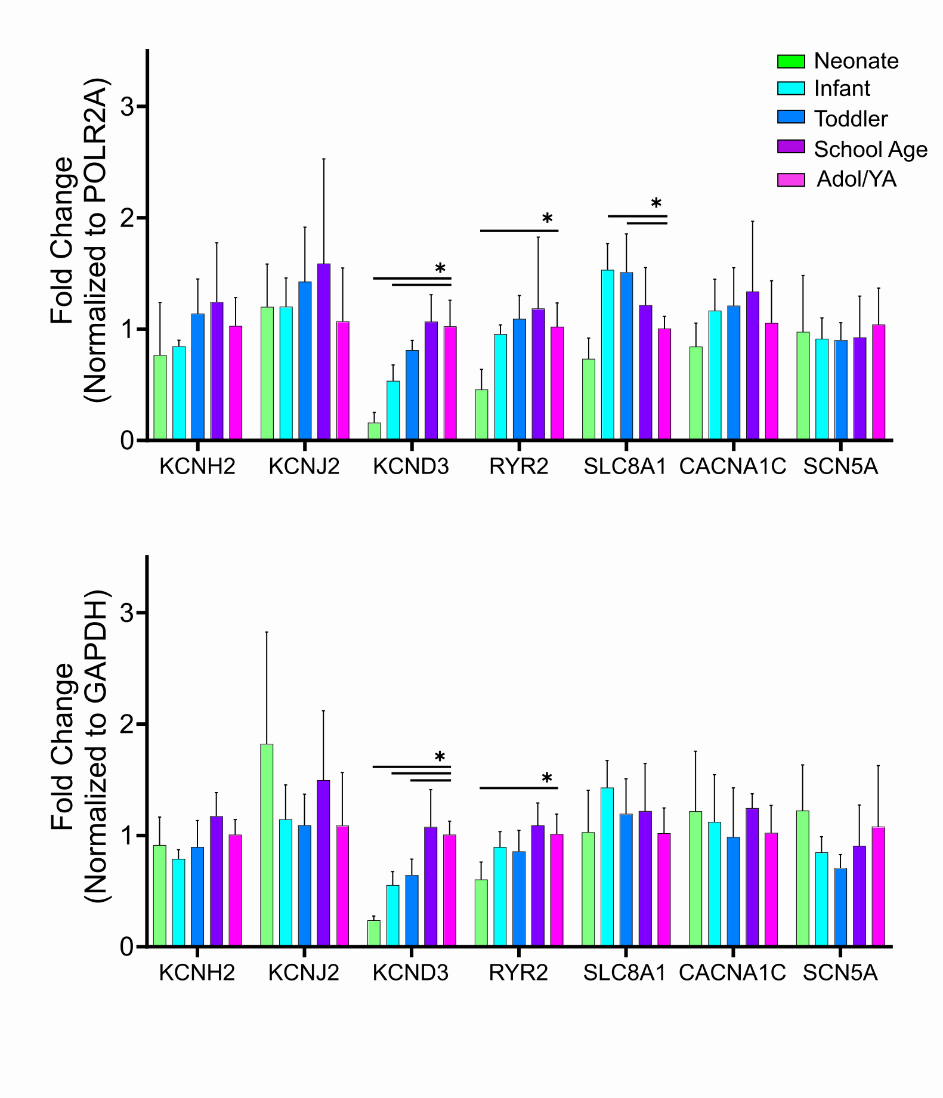
**Supplemental Figure 4.** Real-Time Quantitative Reverse Transcription PCR (qRT-PCR) was used to assess congruency with microarray results shown in Figure 4. A subset of genes and samples were utilized, due to the limited tissue resource material. Genes of interest (*KCNH2, KCNJ2, KCND3, RYR2, SLC8A1, CACNA1C, SCN5A)* were normalized to two housekeeping genes (*GAPDH, POLR2A).* Statistical significance determined by one-way ANOVA with Holm-Sidak test (parametric data) or Kruskal-Wallis with Dunn’s test (non-parametric data). Sample size includes n=3 neonate, n=5 infant, n=5 toddler, n=5 school age, n=6 adolescent/young adults (Adol/YA); *p<0.05 significant difference in fold-change value, relative to oldest age group (Adol/YA).

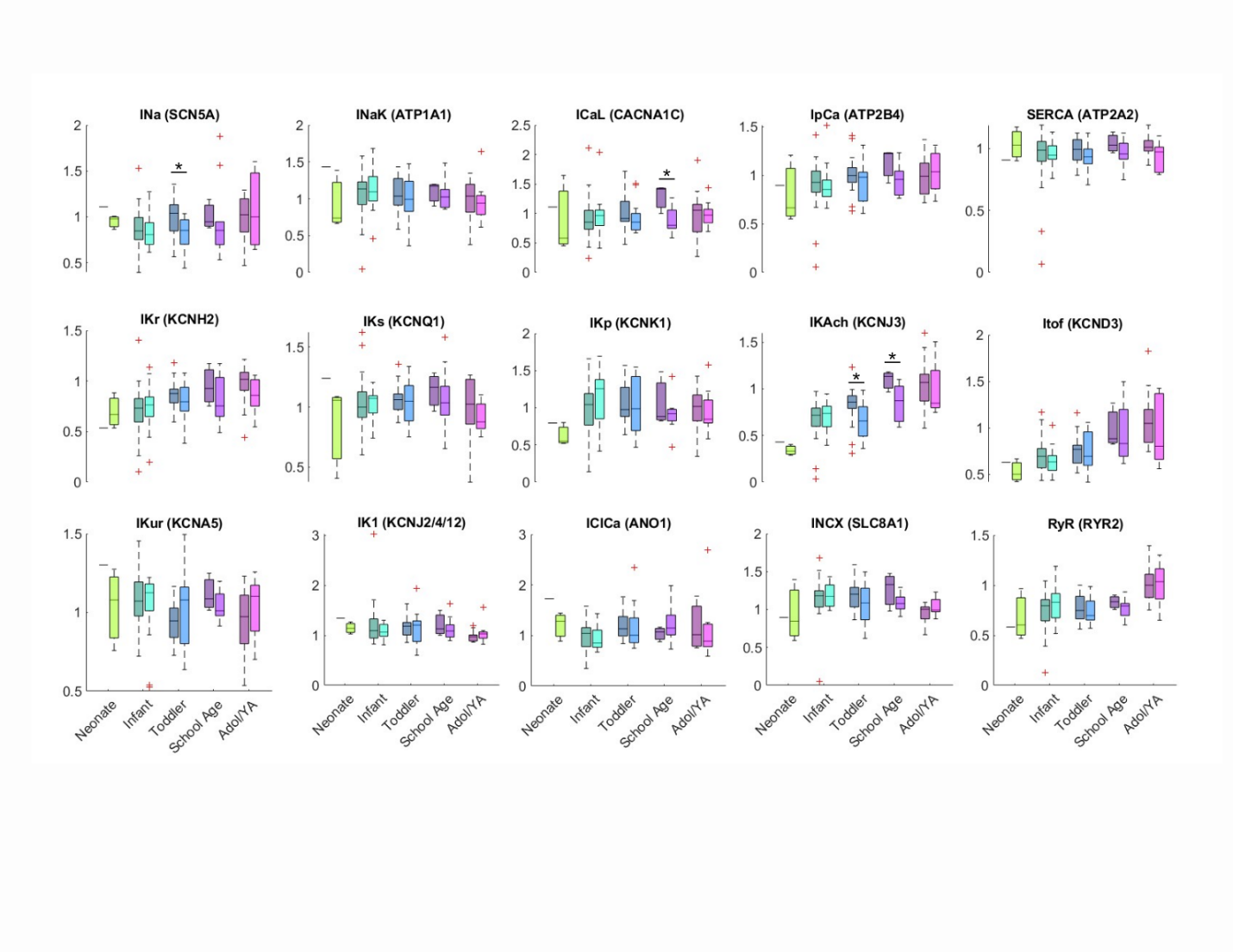

**Supplemental Figure 5.** Sex-specific differences in gene expression within each of the five age groups. Note: units are arbitrary, as raw signals are normalized to the median of the oldest age group (Adol/YA). Within each pair, male patients are represented by the left-most boxplot (darker shade) and females by the right-most boxplot (lighter shade). Adol/YA = adolescents and young adults. Data shown as the median (line), interquartile range (box), statistical outliers (red crosses). Statistical analysis by two-tailed Student’s t-test (parametric) or Kolmogorov-Smirnov test (non-parametric), *p<0.05

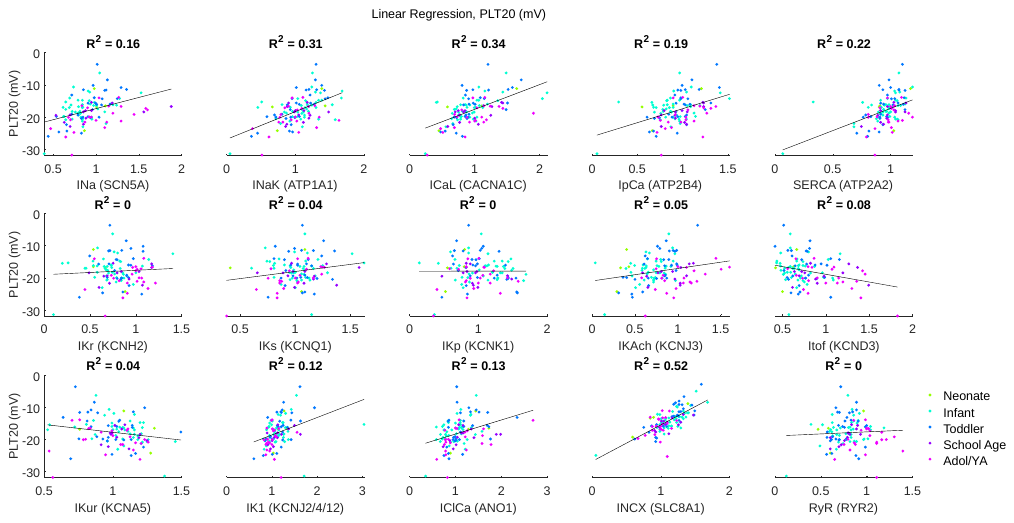

**Supplemental Figure 6.** Quantification of PLT_20_ as a function of each gene included in the model. PLT_20_ was primarily determined by *I*_NCX_ (R^2^ = 0.52); however, *I*_CaL_, *I*_NaK_, and SERCA may also play a significant role in determining the plateau potential (R^2^ = 0.34, 0.31, and 0.22, respectively). Adol/YA = adolescents and young adults.

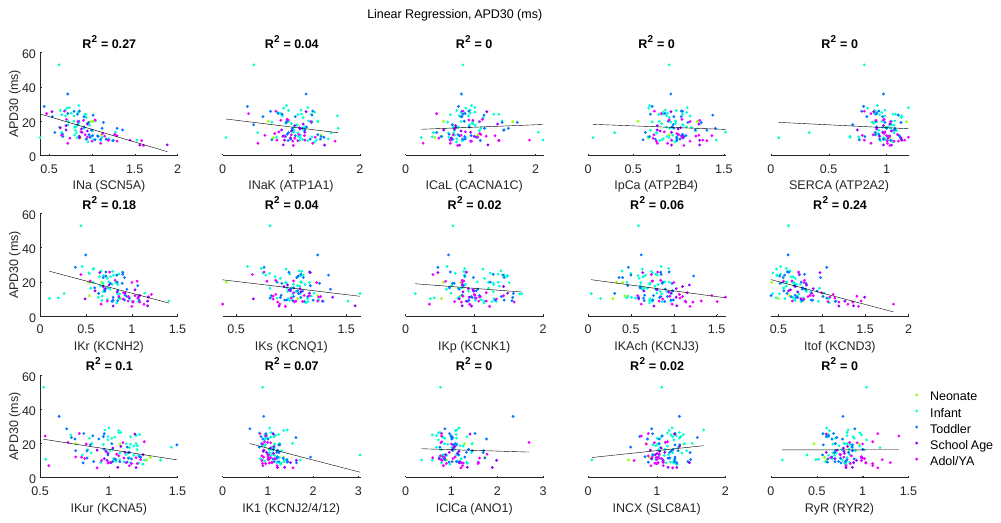

**Supplemental Figure 7.** Quantification of action potential duration at 30% repolarization (APD_30_) as a function of each gene included in the model. Both *I*_Na_ and *I*_tof_ had a correlation greater than 0.2 (R^2^ = 0.27 and 0.24, respectively ), which could contribute to a faster 30% repolarization. Adol/YA = adolescents and young adults.

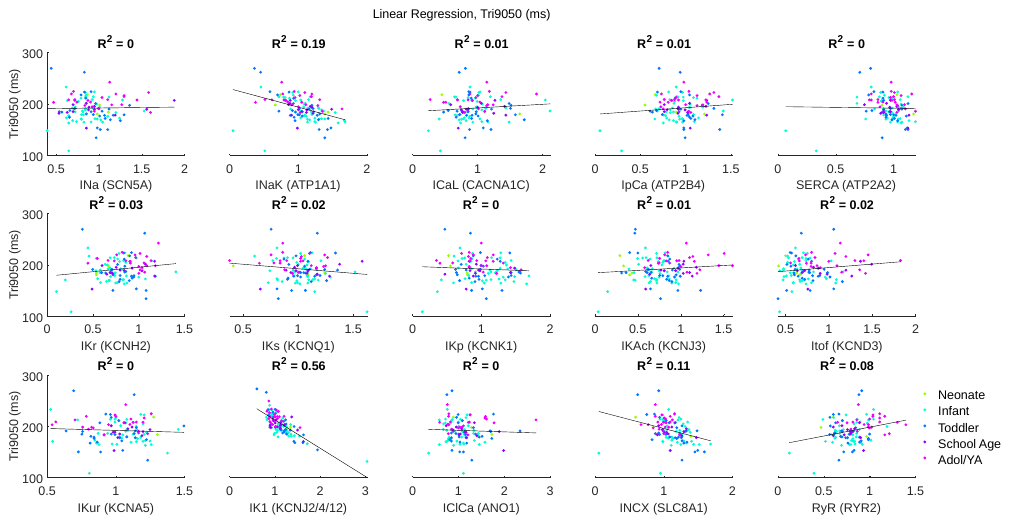

**Supplemental Figure 8.** Quantification of action potential triangulation (Tri_9050_) as a function of each gene included in the model. Only *I*_K1_ had an R^2^ > 0.2 (R^2^ = 0.56), indicating primarily that increases in *I*_K1_ expression lead to a smaller Tri_9050_ and subsequently a less triangular action potential. Adol/YA = adolescents and young adults.

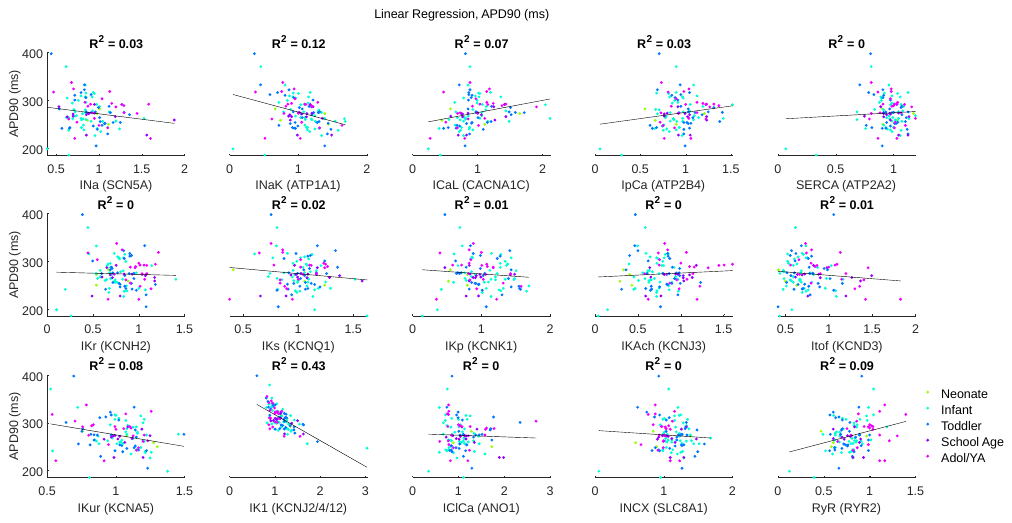

**Supplemental Figure 9.** Quantification of action potential duration at 90% repolarization (APD_90_) as a function of each gene included in the model. Only *I*_K1_ had a correlation greater than 0.2 (R^2^ = 0.43), indicating that increasing *I*_K1_ expression leads to a shorter APD_90_. Adol/YA = adolescents and young adults.

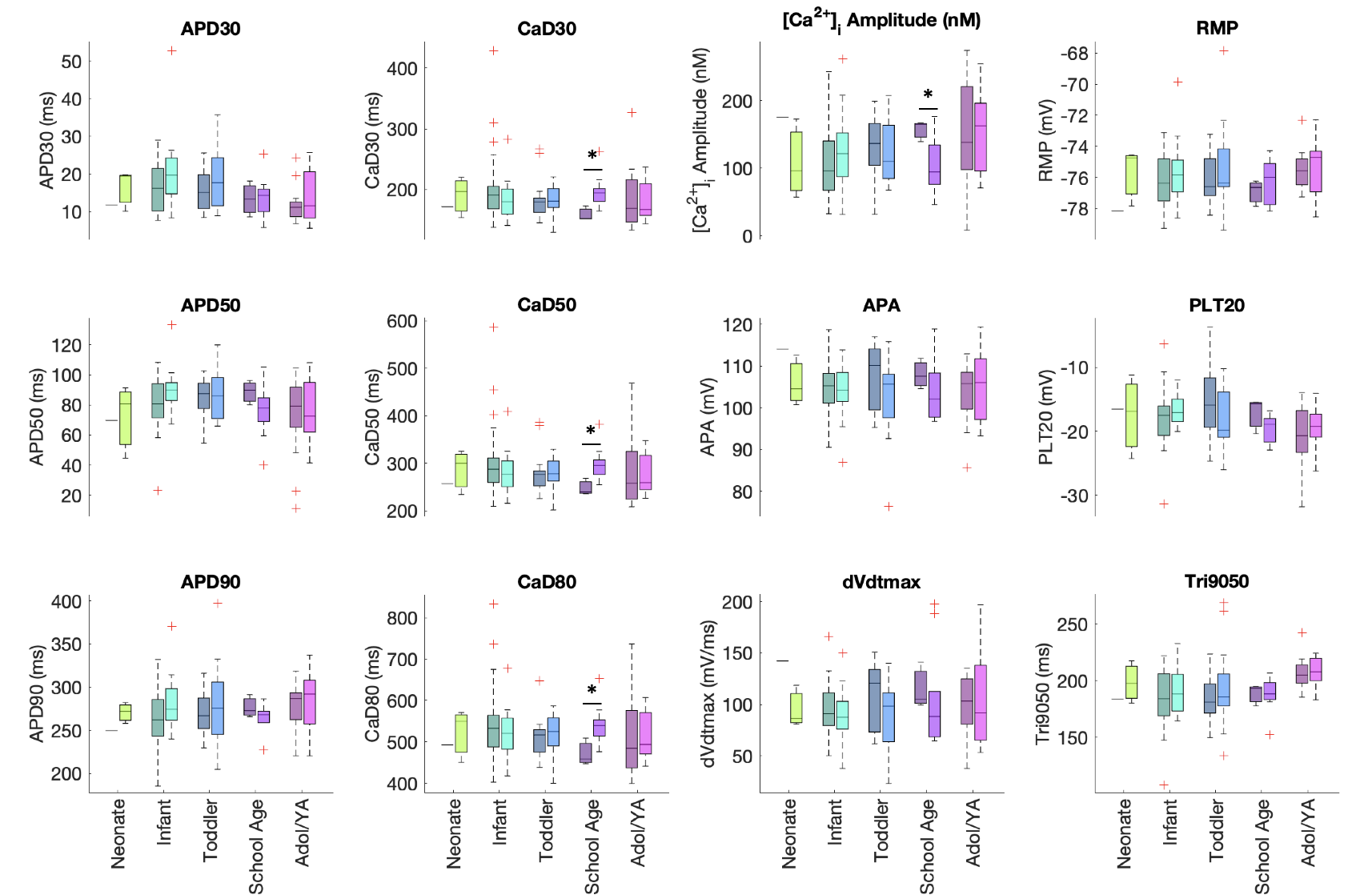

**Supplemental Figure 10.** Sex-based changes in the action potential and calcium transient within each age group, with male patients represented by the left-most boxplot in each pair and illustrated by a darker shade. Significant differences were observed in the calcium transient between male and female patients in the school age group where female patients presented with calcium transients that were significantly longer in duration and shorter in amplitude. *Data shown as the median (line), interquartile range (box), statistical outliers (red crosses).* Statistical analysis by two-tailed Student’s t-test (parametric) or Kolmogorov-Smirnov test (non-parametric), *p<0.05. Abbreviations: adolescents and young adults (Adol/YA), action potential duration at 30% (APD_30_), 50% (APD_50_) and 90% repolarization (APD_90_), plateau potential at 20% of APD_90_ (PLT_20_), action potential amplitude (APA), resting membrane potential (RMP), maximum rate of depolarization (dV/dt_max_), action potential triangulation (Tri_9050_), calcium transient duration at 20% (CaD_20_), 50% (CaD_50_), and 80% (CaD_80_), and the calcium transient amplitude (Ca^2+^ Amplitude).

**Supplemental Table 1.** Patient demographics. Values are reported as median [range] or n-value, with the STAT (Society of Thoracic Surgeons-European Association for Cardio-Thoracic Surgery) score designated using the STAT 2020 category assessment^33^. ASD: atrial septal defect, AVC: atrioventricular canal, COA: coarctation of the aorta, PAPVC: partial anomalous pulmonary venous connection, VSD: ventricular septal defect. Diagnoses in the ‘other’ group included aortic aneurysm, cardiac tumor, and acyanotic tetralogy of Fallot.

| **Age Group** | **Neonate** | **Infant** | **Toddler/**  **Preschool** | **School Age** | **Adolescent/**  **Young Adult** |
| --- | --- | --- | --- | --- | --- |
| No. of Patients | 4 | 46 | 31 | 13 | 23 |
| Age | 18.5 [5-30] days | 135.5 [41-320] days | 2.72 [1.00-5.76] years | 9.26 [6.43-11.98] years | 14.50 [12.04-32.18] years |
| Weight (kg) | 2.6 [2.4-3.0] | 5.1 [3.0-8.3] | 12.5 [7.4-35.5] | 29.4 [16.0-55.6] | 63.0 [22.1-110.0] |
| **Sex** |  | | | | |
| Male | 1 | 26 | 17 | 3 | 14 |
| Female | 3 | 20 | 14 | 10 | 9 |
| **STAT Category** |  | | | | |
| 1 | 1 | 31 | 30 | 1 | 19 |
| 2 | 2 | 13 | 1 | - | 3 |
| 3 | 1 | 1 | - | - | 1 |
| 4 | - | 1 | - | - | - |
| 5 | - | - | - | - | - |
| **Primary Diagnosis** |  | | | | |
| ASD | - | 2 | 17 | 5 | 6 |
| AVC | - | 8 | 1 | - | 1 |
| COA/Aortic Arch Hypoplasia | 3 | 1 | 1 | - | - |
| Coronary Anomaly | - | 1 | - | 2 | 4 |
| PAPVC | - | - | 1 | 2 | - |
| Valve Disease | - | 3 | 3 | 1 | 8 |
| VSD | 1 | 25 | 8 | 3 | 2 |
| Other | - | 6 | - | - | 2 |

**Supplemental Table 2.** Physiological biomarker ranges for action potential and intracellular calcium characteristics in human atrial myocytes. Values were subsequently used as criteria for the generation of physiological cells, based on the qualification that cells had to be within two standard deviations of the mean for all biomarkers (except Ca^2+^ amplitude). Abbreviations: APDxx, AP duration at XX% of repolarization; PLT20, plateau potential at 20% of APD90; APA, action potential amplitude; RMP, resting membrane potential; dV/dt_max_, maximum rate of depolarization; Ca^2+^ amplitude, calcium transient amplitude.

| **Biomarker** | **Mean ± SD** |
| --- | --- |
| APD_90_ (ms) | 317.41 ± 9.33 |
| APD_50_ (ms) | 138.09 ± 9.75 |
| APD_20_ (ms) | 7.22 ± 1.83 |
| PLT_20_ (ms) | -16.28 ± 1.40 |
| APA (mV) | 94.95 ± 1.52 |
| RMP (mV) | -73.98 ± 0.86 |
| dV/dt_max_ (mV/ms) | 219.44 ± 14.65 |
| Ca^2+^ amplitude | > 50 nM |

| **Ionic Current** | **Associated Gene** | **Source** |
| --- | --- | --- |
| INa | SCN5A | Smirnov et al. |
| INab | Unknown | n/a |
| INaK | ATP1A1 | Smirnov et al. |
| ICaL | CACNA1C | Smirnov et al. |
| ICab | Unknown | n/a |
| IpCa | ATP2B4 | Smirnov et al. |
| SERCA | ATP2A2 | Smirnov et al. |
| IKr | KCNH2 | Smirnov et al. |
| IKs | KCNQ1 | Smirnov et al. |
| IKp | KCNK1 | GenBank |
| IKAch | KCNJ3 | GenBank |
| Itof | KCND3 | Wang et al. |
| Ikur | KCNA5 | GenBank |
| IK1 | KCNJ2/KCNJ4/KCNJ12 | Reilly and Eckhart (see text for details) |
| IClCa | ANO1 | GenBank |
| IClb | Unknown | n/a |
| INCX | SLC8A1 | Smirnov et al. |
| RyR | RYR2 | Smirnov et al. |

**Supplemental Table 3.** Gene selection was initially based on published data by Smirnov et al, with the following modifications for atrial cells: KCND3 was used to model *I*_tof_, an average of all isoforms (KCNJ2, KCNJ4, KCNJ12) was used to model *I*_K1._ The associated gene for any additional atrial-specific currents was identified using the NIH tool GenBank.

**Supplemental Table 4.** Patient arrhythmia prediction based on simulated arrhythmia score. Results shown as area under the curve (AUC) from receiver operating characteristics (ROC) curve. Patient arrhythmia data documentation included n=86 preoperative assessment, and n=90 postoperative assessment. Sample size indicates the number of individuals presenting with an arrhythmia, out of the total number of patients with data documentation. Some patients presented with multiple arrhythmia sub-types, and as such, the sub-type sum will not always be equal to the total arrhythmia type sum (e.g., preoperative premature atrial contractions + miscellaneous ectopy = 5, preoperative total ectopy = 4). Abbreviations: atrioventricular (AV), bundle branch block (BBB), conduction delay (CD), sinoatrial (SA).

| **Preop group**  **(n = 86)** | **Preop subgroup** | **AUC** | **n** | **Postop group**  **(n = 90)** | **Postop subgroup** | **AUC** | **n** |
| --- | --- | --- | --- | --- | --- | --- | --- |
| Any Preop Arrhythmia |  | 0.544 | 54 | Any Postop Arrhythmia |  | 0.550 | 87 |
| BBB/CD |  | 0.533 | 15 | BBB/CD |  | 0.667 | 42 |
| QRST Abnormality |  | 0.542 | 43 | QRST Abnormality |  | 0.360 | 72 |
| Sinus/Atrial Arrhythmias | All | 0.436 | 20 | Sinus/Atrial Arrhythmias | All | 0.563 | 47 |
|  | PR Abnormality | N/A | 0 |  | PR Abnormality | 0.439 | 6 |
|  | Sinus Node Dysfunction/Sinus Pause | N/A | 0 |  | Sinus Node Dysfunction/Sinus Pause | 0.405 | 5 |
|  | Slow Junctional/Junctional Escape | N/A | 0 |  | Slow Junctional/Junctional Escape | 0.508 | 8 |
|  | AV Block | 0.480 | 4 |  | AV Block | 0.652 | 11 |
|  | Sinus Bradycardia | 0.445 | 4 |  | Sinus Bradycardia | 0.533 | 13 |
|  | Sinus Tachycardia | 0.547 | 7 |  | Sinus Tachycardia | 0.503 | 17 |
|  | Miscellaneous SA Arrhythmias | 0.278 | 5 |  | Miscellaneous SA Arrhythmias | 0.595 | 23 |
| Ectopy | All | 0.378 | 4 | Ectopy | All | 0.447 | 18 |
|  | Premature Atrial Contractions | 0.673 | 2 |  | Premature Atrial Contractions | 0.610 | 12 |
|  | Miscellaneous Ectopy | 0.225 | 3 |  | Miscellaneous Ectopy | 0.343 | 9 |

**Supplemental Table 5.** Differentially expressed genes, as compared to the oldest patient group (adolescent/young adults) using a 1.25-fold cut off and a false discovery rate of 0.1.

**Supplemental Table 6.** Unique annotation clusters in neonates or infants, as compared to the oldest patient group (adolescent/young adults). Analysis performed using DAVID, rank-ordered by p-value (cut-off using Bonferroni-adjusted p-value of 0.1).

**Supplemental Table 7.** Enriched canonical pathways in neonates or infants, as compared to the oldest patient group (adolescent/young adults). Analysis performed using Ingenuity Pathway Analysis, rank-ordered by p-value (cut-off using Fisher’s exact test with adjusted p-value of 0.1).

**Supplemental Table 8.** Enriched gene ontologies in neonates or infants, as compared to the oldest patient group (adolescent/young adults). Analysis performed using Enrichr, rank-ordered by p-value (cut-off using adjusted p-value of 0.2).
